# A geometry-over-coevolution principle governs protein complex assembly in AlphaFold

**DOI:** 10.64898/2026.04.03.716280

**Authors:** Shuangjun Li, Zichun Mu, Chengfei Yan

## Abstract

AlphaFold has revolutionized protein complex structure prediction, yet how it assembles intermolecular interfaces remains poorly understood. Contrary to the prevailing view that inter-protein coevolution drives complex prediction, we uncover a geometry-over-coevolution principle governing assembly in AlphaFold-Multimer and AlphaFold3. Through systematic perturbation of evolutionary and structural inputs and development of residue-level constraint propagation mapping to trace the emergence and propagation of geometric information within the network, we show that prediction accuracy is governed primarily by monomer-derived structural geometry and interface-specific sequence–geometry compatibility, rather than direct inter-protein coevolutionary signals. Our mapping reveals a hierarchical assembly mechanism in which monomer-level geometric representations are established first and progressively propagated to constrain cross-chain interfaces. This mechanism further explains why antigen–antibody complexes are predicted less accurately, as their intrinsic interface plasticity and non-canonical architectures limit the propagation of geometric constraints across interfaces. Together, these findings establish a mechanistic framework for understanding how artificial intelligence models assemble protein complexes and provide principles for improving next-generation structure prediction.

## Introduction

The physical organization of cellular systems fundamentally relies on the precise assembly of protein-protein complexes. While determining their three-dimensional architectures has long been a central challenge in structural biology, recent advances in artificial intelligence (AI) —most notably the AlphaFold family of models^1–3^, including AlphaFold-Multimer (AFM)^2^ and AlphaFold3 (AF3)^3^, have revolutionized this field by achieving unprecedented predictive accuracy. However, this empirical success has drastically outpaced our fundamental understanding of the models themselves; the mechanistic principles by which these deep learning architectures accurately recognize and assemble multi-chain interfaces remain largely treated as a “black box”. Uncovering these internal mechanisms is not merely an exercise in AI interpretability; it is essential for understanding the physical determinants of prediction accuracy and for guiding the development of more accurate and mechanistically informed models of biomolecular interactions.

A prevailing paradigm attributes AlphaFold’s success in complex prediction primarily to inter-protein coevolutionary information encoded within multiple sequence alignments (MSAs)^2–11^. This assumption heavily extrapolates from monomeric structure prediction, where intra-protein correlated mutations provide robust evolutionary constraints for defining residue-residue contacts^1,3,4,12^. Consequently, the field has widely presumed that inter-protein coevolution provides analogous spatial constraints for mapping intermolecular interfaces^13–19^. Yet, this coevolution-centric view is increasingly contradicted by empirical observations: inter-protein coevolution is often drastically weaker or entirely absent—such as in transient interactions and structurally immune recognition recognition—where AlphaFold nevertheless frequently produces plausible complex structures^8,10,20–24^. This apparent paradox challenges the prevailing view and raises a fundamental mechanistic question: how does AlphaFold accurately assemble protein complexes in the absence of strong inter-protein coevolutionary information?

Here, we resolve this paradox by systematically dissecting the mechanistic basis of complex assembly in AFM and AF3 through controlled perturbations of evolutionary and structural inputs. We demonstrate that inter-protein coevolution makes, at best, a marginal contribution to accurate complex prediction. Instead, we reveal that protein-protein interfaces are governed predominantly by monomer-derived geometric constraints coupled with localized sequence-geometry compatibility.

To explicitly visualize this hidden inference process, we develop an interpretability framework, AlphaFold-Constraint Propagation Mapping (AF-CPM). By extracting and mapping latent pair representations across Evoformer layers and successive recycling iterations, AF-CPM allows us to dynamically trace the spatiotemporal evolution of geometric constraints at the residue level. This tracking uncovers a hierarchical assembly mechanism: robust monomeric structural geometries crystallize well before the emergence of inter-chain contacts. Consequently, AlphaFold does not “read” inter-chain interfaces directly from paired evolutionary sequences; rather, it progressively “builds” them by propagating pre-established monomeric constraints to refine structural complementarity

Finally, we leverage this geometry-driven framework to interrogate one of the most notoriously challenging targets for current AI models: antigen-antibody complexes^3,8,20,24,25^. We show that the reduced predictive accuracy in immune recognition is not a consequence of absent coevolutionary signals, but rather stems from the profound structural plasticity and statistically atypical architectures of immune interfaces, which hinder the propagation of geometric constraints needed for accurate interface assembly. Together, our findings reveal a geometry-over-coevolution principle underpinning AlphaFold-based complex assembly. By replacing the coevolution-centric dogma with a geometry-grounded mechanistic framework, this work provides a conceptual foundation for decoding AI behavior in structural biology and guiding the development of next-generation models optimized for highly flexible and non-canonical biomolecular interactions.

## Results

### 1. Overview of the modeling frameworks of AFM and AF3

We begin by outlining the modeling pipelines of AFM and AF3 for protein complex structure prediction. Both frameworks take amino acid sequences and their corresponding MSAs as inputs, optionally incorporating monomeric structural templates. Following input embedding, representations are processed by the Evoformer in AFM or the Pairformer in AF3, where intra- and inter-protein geometric relationships are jointly modeled. These representations are then passed to the structure generation module, which is built on Invariant Point Attention (IPA) in AFM and a diffusion-based generative model in AF3, to generate the three-dimensional structure of the protein complex^2,3^ (Figure 1a).

**Figure 1.**
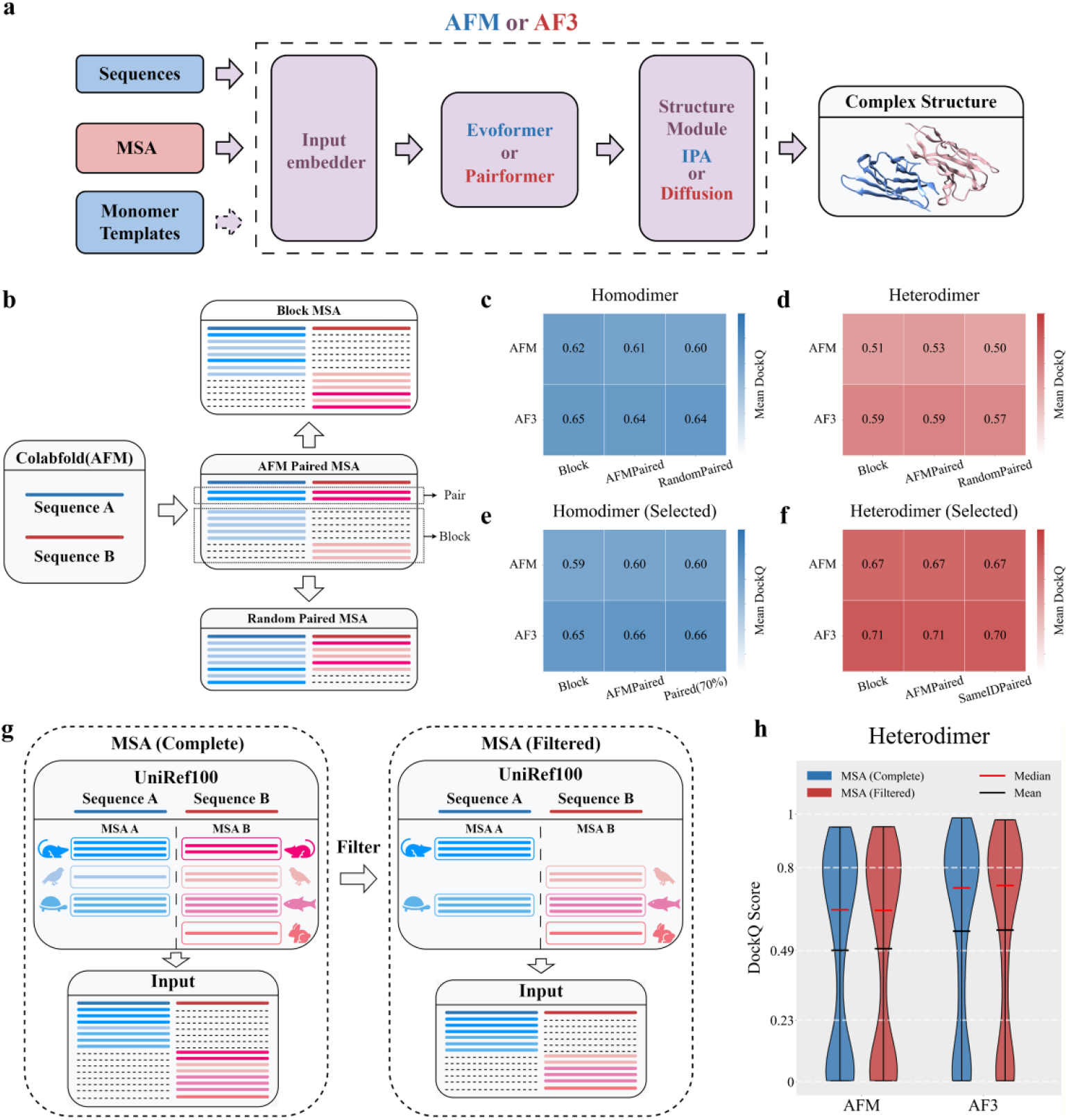
Effects of MSA pairing strategies on protein complex structure prediction. (a) AFM and AF3 pipelines for protein complex modelling. (b) Schematic of MSA input strategies: Block MSA, AFM Paired MSA, and Randomly Paired MSA. (c, d) Mean DockQ scores for homodimer (c) and heterodimer (d) datasets using AFM and AF3 with different MSA inputs. (e) Performance on the filtered homodimer dataset using Block MSA, AFM Paired MSA, and Block MSA with paired sequences at ≥70% identity to the target. (f) Performance on the filtered heterodimer dataset using Block MSA alone, AFM Paired MSA, and Block MSA with paired sequences sharing identical UniProt IDs. (g) Schematic of Block MSA construction in which the MSAs of the two interacting proteins contain no shared species. (h) Performance of AFM and AF3 with or without species overlap between partner MSAs. The white dashed horizontal line represents key thresholds based on the DockQ evaluation of the predicted results: incorrect (0 ∼ 0.23), acceptable quality (0.23 ∼ 0.49), medium quality (0.49 ∼ 0.8), and high quality (0.8 ∼ 1).

Both models employ phylogeny-aware MSA pairing strategies^26^ intended to capture putative inter-protein coevolutionary signals that may inform interface formation. However, the contribution of inter-protein coevolution to accurate complex structure prediction, whether explicitly encoded through paired MSAs or implicitly embedded in unpaired MSAs, remains unclear and warrants systematic investigation.

### 2. Impact of inter-protein coevolutionary information on AFM and AF3 predictions

#### 2.1 Effect of paired MSA-derived inter-protein coevolution on prediction accuracy

To rigorously assess the contribution of paired MSAs to protein complex structure prediction, we constructed a strictly time-segregated benchmark comprising Protein Data Bank (PDB)^27^ entries deposited between January 1, 2022 and December 20, 2024, excluding all structures present in the training sets of AFM and AF3 (cutoff: September 30, 2021). After applying resolution and redundancy filters, the final dataset included 200 homodimers and 316 heterodimers. Homodimers were analyzed separately, as their paired MSAs are generated by concatenating identical sequences without cross-chain pairing, avoiding confounding effects from erroneous pairings.

Using ColabFold^28^, we first generated the native AFM Paired MSAs^2^ for each protein pair, which consist of (i) paired sequences derived from phylogenetically matched homologs and (ii) unpaired sequences organized in a block-diagonal format to preserve alignment consistency. For homomeric targets, this construction naturally reduces to the concatenation of identical monomeric sequences, thereby eliminating the block-diagonal component.

To isolate the effect of sequence pairing, we constructed two controlled variants from the same underlying MSAs: Block MSA, in which all non-query sequences were gap-padded to eliminate inter-chain pairing, and Randomly Paired MSA, in which sequences were randomly paired across chains. These three conditions form a controlled framework for disentangling the role of paired MSAs (Figure 1b). Each alignment was input to AFM and AF3 without templates, and performance was evaluated using DockQ^29^ across both homo- and heteromeric complexes.

As shown in Figures 1c-d, model performance measured by mean DockQ shows minimal sensitivity to the type of MSA. Predictions from Block MSAs and native AFM MSAs are nearly indistinguishable (|Δ| ≤ 0.2), indicating that explicit sequence pairing contributes little to predictive accuracy. Randomly Paired MSAs yield only a marginal decrease (Δ ∈ [−0.03, 0]), likely due to noise introduced by incorrect pairings. To exclude potential bias from training-set similarity, we further evaluated a subset of complexes with low structural similarity to the AFM and AF3 training data (Supplementary Text S1, Figure S1), thereby minimizing reliance on template-based recognition. Performance on this subset remains consistent with that on the full dataset, with no significant differences across the three MSA strategies. Collectively, these results indicate that inter-protein coevolutionary signals encoded in paired MSAs are not a major determinant of complex structure prediction accuracy in either AFM or AF3.

Although introducing paired MSAs does not result in a significant difference in overall prediction performance (mean DockQ), a case-by-case analysis of predictions from AFM MSA and Block MSA reveals that paired MSAs can indeed improve prediction accuracy in some individual cases (Supplementary Figure S2). However, there are also instances where the introduction of paired MSAs leads to a decrease in prediction accuracy. Therefore, the limited benefits of AFM-generated paired MSAs may stem, in part, from imperfections in the pairing strategy itself. For homomeric interactions, AFM concatenates homologous sequences irrespective of whether they share the same oligomerization mode, such that low-similarity sequences may introduce noise. For heteromeric interactions, AFM’s phylogeny-guided pairing does not guarantee that concatenated sequences correspond to bona fide interacting partners.

Notably, in the cases where paired MSAs significantly improve prediction accuracy, the overlap between the AFM and AF3 model predictions is only 20–30% (Supplementary Figure S2). This raises the question of whether the observed improvement is driven by the coevolutionary information encoded in the paired MSAs, or if it results from the inherent randomness and biases of the models. If incorrect pairings are minimized, can paired MSAs truly enhance the overall performance of the models?

To better assess the ideal contribution of paired MSAs to prediction performance and minimize the impact of incorrect pairings, we applied additional stringency filters. For homodimers, only paired sequences with more than 70% sequence identity to the target were retained, while lower-identity homologs were reassigned to the unpaired MSA. For heterodimers, only sequence pairs sharing the same UniProt^30^ ID were preserved, while all other sequences were placed in the unpaired component. Sequence pairs with identical UniProt IDs unambiguously originate from the same protein and typically correspond to distinct domains; such pairs therefore represent correct pairings that do not introduce noise from non-interacting proteins. After filtering, we retained interactions containing more than 50 paired sequences, yielding 61 homomeric and 54 heteromeric complexes for subsequent modeling with AFM and AF3. Despite this stringent selection, no significant performance improvement was observed (Figures 1e–f), reinforcing the conclusion that inter-protein coevolutionary signals in paired MSAs are not a primary driver of AFM or AF3 performance in protein complex structure prediction.

#### 2.2 Effect of latent inter-protein coevolution in unpaired MSAs on prediction accuracy

The lack of performance gains from paired MSAs does not, by itself, exclude a role for inter-protein coevolutionary signals in complex structure prediction. One possibility is that, during training, the models may have implicitly learned interaction patterns across large collections of unpaired sequences, thereby acquiring the capacity to extract inter-protein coevolutionary information even in the absence of explicit sequence pairing.

To directly test whether AFM and AF3 rely on inter-protein coevolutionary signals potentially embedded within unpaired MSAs when predicting complex structures, we sought to eliminate any such signals to evaluate the resulting impact on model performance. This task is nontrivial, as AFM constructs MSAs from multiple metagenomic databases that contain numerous sequences lacking reliable species annotations, making it difficult to ensure that unpaired MSAs do not include sequences derived from the same organism and thus retain latent species-level coevolutionary information.

To overcome this limitation, we regenerated MSAs for proteins in the heterodimer dataset using JACKHMMER^31^ searches against the UniRef100^32^ database, retaining only sequences with explicit species annotations. These monomeric MSAs were concatenated in block-diagonal format to generate Block MSAs. To further eliminate any inter-protein coevolutionary signal embedded in Block MSAs, we randomly selected two non-overlapping sets of species for the two monomer MSAs and retained only sequences originating from their respective species sets. This procedure ensured that the two MSAs shared no common species, thereby removing potential species-level interaction signals. Block-format interaction MSAs were then reconstructed using the filtered sequences (Figure 1g). These two sets of Block MSAs were subsequently provided as input to AFM and AF3 for heterodimeric complex structure prediction. Model performance showed no significant difference between the two conditions (Figure 1h), indicating that latent inter-protein coevolutionary signals potentially embedded in unpaired MSAs are not a major contributor to predictive accuracy.

Taken together, these results demonstrate that inter-protein coevolutionary information, whether explicitly encoded through paired MSAs or implicitly embedded within unpaired MSAs, is not the primary determinant of AFM or AF3 performance in protein complex structure prediction.

### 3. Mechanism of complex prediction without inter-protein coevolution

#### 3.1 A hypothesis: Inter-protein spatial constraints derived from monomer geometries

Our finding that inter-protein coevolution is not the primary source of information used by AFM and AF3 raises a central question: how do these models nonetheless achieve high accuracy in protein complex prediction?

A theoretical framework proposed by Roney *et al.*^33^ for protein monomer structure prediction posits that AlphaFold2 exploits coevolutionary signals encoded in MSAs to guide the model toward regions near the global minimum of the free-energy landscape, followed by local refinement to reach the lowest-energy state, with model confidence metrics reflecting this underlying landscape (Figure 2a).

**Figure 2.**
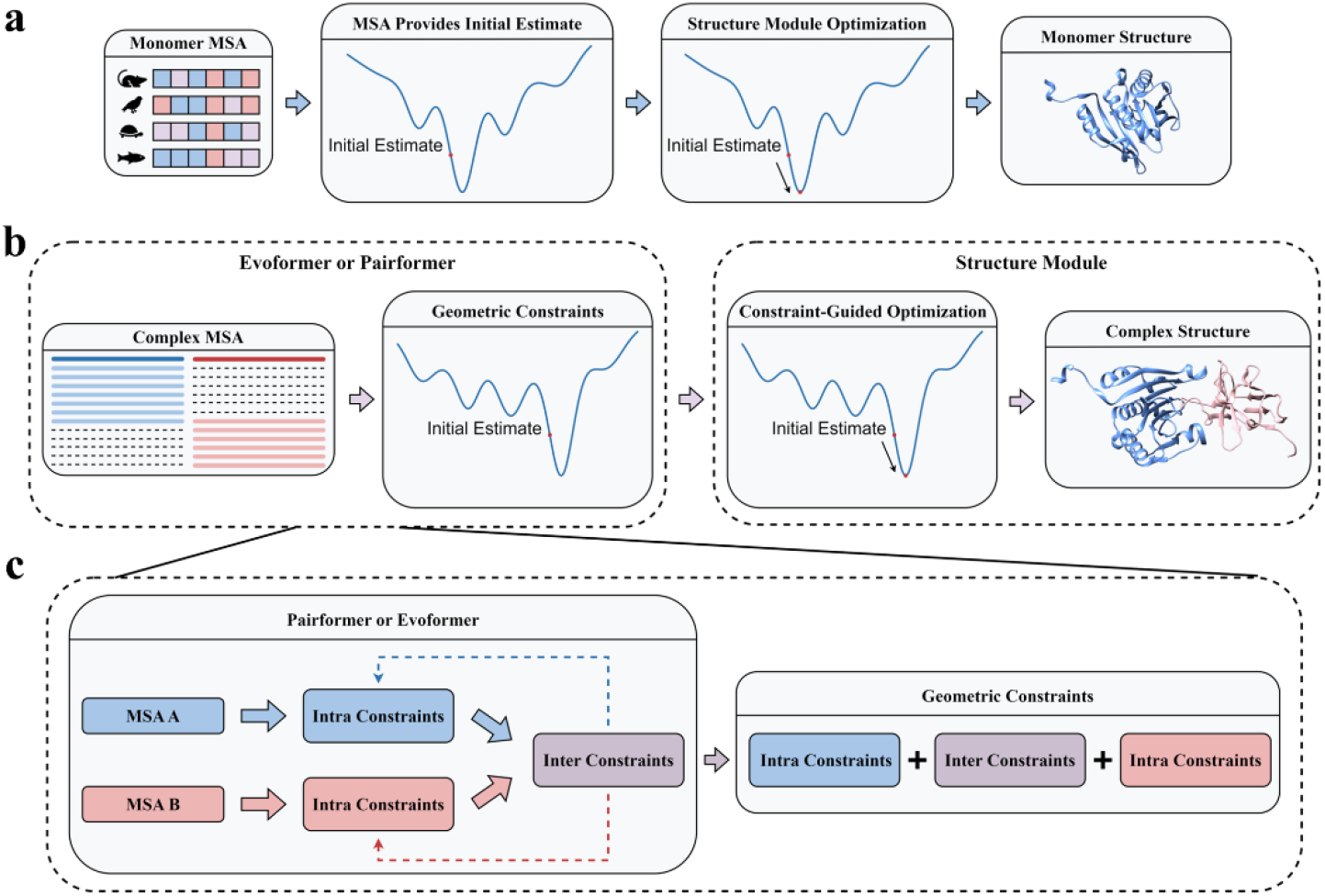
Mechanistic insights into AlphaFold predictions of monomeric and complex structures from MSAs. (a) AlphaFold generates an initial structural estimate from monomer MSAs, which is refined by the structure module to yield the final monomer structure. Image inspired by Roney et al.^33^. (b) For protein–protein complexes, geometric constraints inferred from complex MSAs through the Evoformer/Pairformer module provide an initial estimate of the complex structure, which is subsequently refined in the structure module to yield the final complex structure. (c) In the Evoformer/Pairformer module, intra-chain constraints are first extracted from monomer MSAs and used to initialize inter-chain constraints, which are then progressively refined through iterative updates, yielding a self-consistent representation for complex structure prediction.

Here, we extend this framework to protein complexes to describe how information is integrated across interacting chains. In AFM and AF3, the representation-learning modules (Evoformer and Pairformer) transform MSA-derived signals into geometric constraints that restrict the conformational search space, guiding the model toward regions near the free-energy minimum. The structure module then refines these candidates toward the lowest-energy state, while confidence metrics continue to reflect the underlying energy landscape (Figure 2b).

This perspective motivates a closer examination of the nature and origin of the geometric constraints governing protein–protein interactions. These constraints can be decomposed into intra-monomer and inter-protein components. While intra-monomer constraints are largely informed by coevolutionary signals within monomer MSAs, the origin of inter-protein constraints remains unclear. Our results indicate that inter-protein coevolution plays a negligible role in complex prediction, suggesting that these constraints are not directly encoded by such signals. Instead, given that binding interfaces are fundamentally determined by the three-dimensional architectures of the individual subunits, we propose that inter-protein spatial constraints are primarily inferred from intrinsic monomer geometries. Specifically, given a protein complex MSA (either paired or block-formatted), the model first infers intra-chain geometric constraints from the monomer sequence information it contains; on this basis, it derives initial inter-chain constraints. These are then iteratively refined through multiple Evoformer/Pairformer layers and recycling rounds, enabling the joint optimization of intra- and inter-chain geometries and ultimately yielding a self-consistent representation for complex structure modeling (Figure 2c).

#### 3.2 Interrogating the hypothesis through template-driven complex prediction

Building on our hypothesis that inter-protein spatial constraints are primarily inferred from monomer geometry rather than direct coevolution, we reasoned that monomer templates explicitly encoding accurate monomer geometry should be sufficient to support complex structure prediction (Figure 3a). Although AFM and AF3 allow the use of monomeric templates, these are generally treated as auxiliary to MSAs, and their standalone contribution remains unclear.

**Figure 3.**
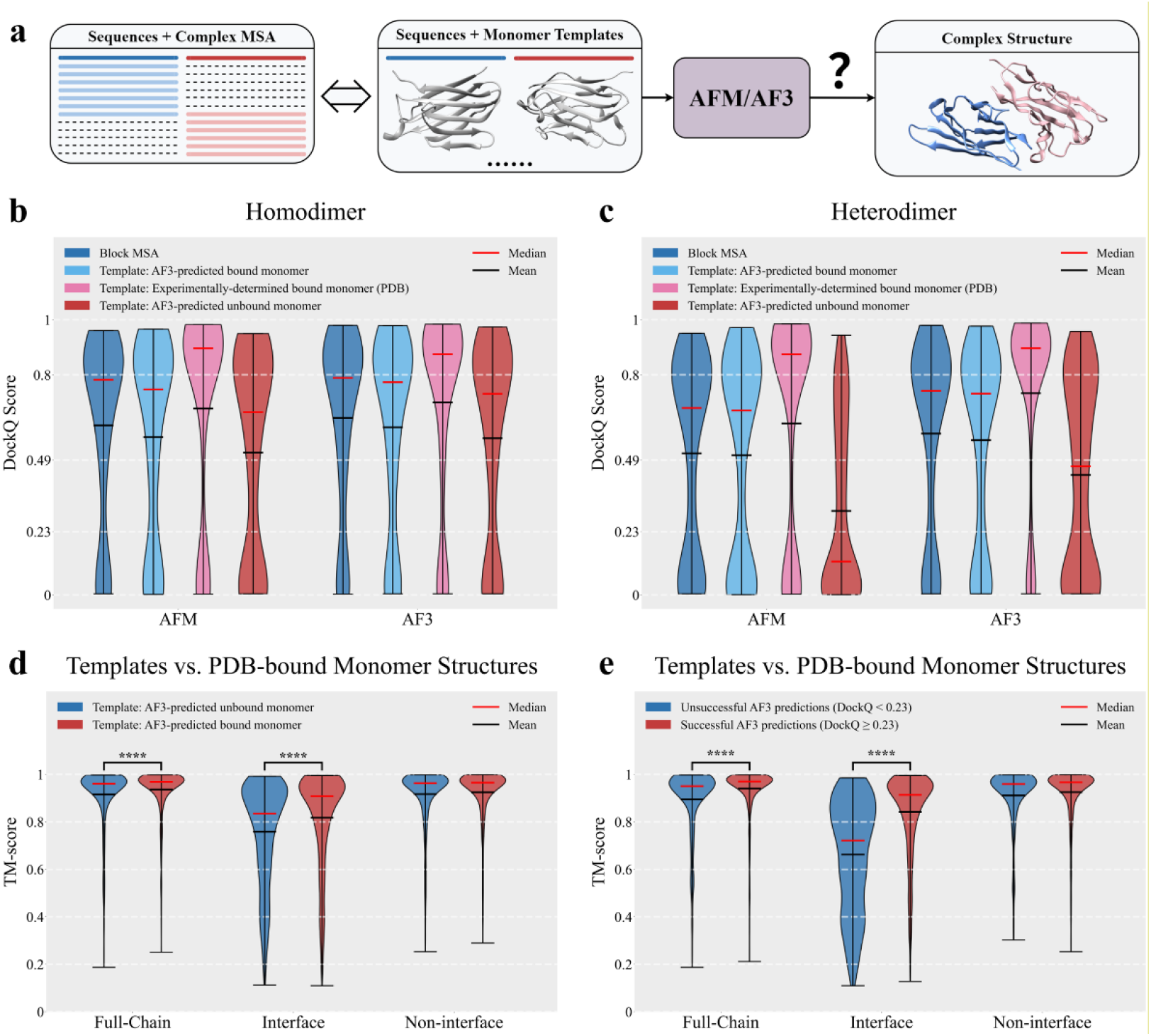
Monomer template-driven protein complex structure prediction. (a) Use of monomer templates as substitutes for complex MSA in complex prediction. (b, c) Performance on homodimer (b) and heterodimer (c) datasets using Block MSA versus three types of monomer templates: AF3-predicted bound templates, experimentally determined bound monomer templates (PDB), and AF3-predicted unbound templates. (d) TM-score distributions of predicted bound and unbound monomer templates relative to experimental bound monomers (PDB), evaluated at the full-chain, interface, and non-interface regions. Significant differences were observed at the full-chain and interface regions (\*\*\*\**p*< 0.0001, two-sided Mann–Whitney U test), but not at the non-interface region (*p*>0.05). (e) TM-score distributions of predicted monomer templates relative to experimental bound monomers (PDB), comparing successful and unsuccessful AF3 predictions at the full-chain, interface, and non-interface regions. Significant differences were observed at the full-chain and interface regions (\*\*\*\**p*<0.0001), but not at the non-interface region (*p* >0.05).

To test this, we evaluated AFM and AF3 using only target sequences and monomer templates. We considered three template types: bound-state monomers derived from AF3 predictions under Block MSA conditions, experimentally determined bound-state monomers, and unbound-state monomers predicted independently by AF3. Performance was assessed on curated homo- and heterodimer datasets and compared to predictions using Block MSAs.

Consistent with our hypothesis, predicted bound-state templates achieved accuracy comparable to MSA-based predictions, indicating that monomer geometry captures much of the structural signal for complex assembly typically attributed to evolutionary information. Notably, experimentally determined bound-state templates even yielded substantially higher accuracy, and incorporating MSAs in addition to such templates did not further improve performance (Figure 3b–c; Supplementary Figure S3). These results demonstrate that accurate protein complex structures can be inferred in the absence of MSAs when high-quality monomer geometry is available. However, predicted unbound-state templates lead to a marked reduction in accuracy, highlighting a clear difference between bound- and unbound-state templates that substantially impacts model performance.

To identify the structural determinants of this performance difference, we compared predicted bound and unbound monomer templates to experimentally determined bound conformations using TM-score^34^ at the full-chain, interface, and non-interface regions. Predicted bound monomer templates more closely resemble experimental conformations, with improvements largely confined to interface regions, while non-interface regions show no significant differences (Figure 3d). Further analysis of AF3 predictions using both bound-state and unbound-state templates revealed that successful complex predictions (DockQ ≥ 0.23) are associated with significantly higher template accuracy at the interface, whereas no such difference is observed in non-interface regions (Figure 3e). Consistent with this observation, interface template accuracy shows a significant correlation (Pearson *r* = 0.575) with complex prediction performance (Supplementary Figure S4).

Collectively, these results indicate that the accuracy of interface regions in monomer templates is a key determinant of complex structure prediction performance, implying that the model may leverage geometric complementarity at protein interfaces to infer inter-chain constraints and guide complex assembly. Notably, interface regions of bound-state monomer structures predicted by AF3 using Block MSA as input exhibit substantially higher accuracy than those of unbound-state monomers predicted using monomer MSAs. This difference propagates to complex structure prediction when these structures are used as monomer templates by AFM/AF3, resulting in deviations in both the global binding mode (Figure 3b–c) and the local interface geometry (Supplementary Figure S5) of the predicted complexes. Together, these observations also suggest that MSA-based modeling of protein–protein complexes allows the model to partially capture interface rearrangements induced by protein–protein interactions, whereas template-only inference has limited ability to correct such discrepancies when the interface geometry deviates from the experimentally observed conformation. Consequently, monomer templates cannot fully replace MSAs in practical applications.

#### 3.3 Effects of interface and non-interface mutations on complex structure prediction

The TM-score-based analysis above primarily evaluates backbone conformations and does not explicitly capture the contribution of side chains—the sequence itself—to protein–protein complex prediction. Consequently, it remains unclear whether interface recognition is governed solely by backbone geometric constraints or also depends on side-chain identity.

To systematically assess the role of side chains, we introduced random mutations separately at interface and non-interface residues in both homomeric and heteromeric protein–protein interaction datasets, with mutations limited to ≤10% of residues per chain. The resulting mutants were evaluated using two complementary prediction strategies: (i) Block MSA-based prediction, in which MSAs were reconstructed from the mutated sequences, and (ii) monomer template-based prediction, in which structural templates were remodeled to reflect the mutations based on experimentally determined bound-state conformations. By comparing the predicted complexes with their corresponding experimental structures, we quantified the impact of side-chain perturbations on complex assembly under distinct structural and evolutionary information regimes.

Strikingly, mutations at interface residues led to a near-complete loss of predictive accuracy under both settings, whereas mutations at non-interface residues had only a moderate effect (Figure 4a–d). These results demonstrate that side-chain identity at protein–protein interfaces is essential for accurate identification of interaction interfaces and prediction of complex structures, extending beyond backbone geometric complementarity.

**Figure 4.**
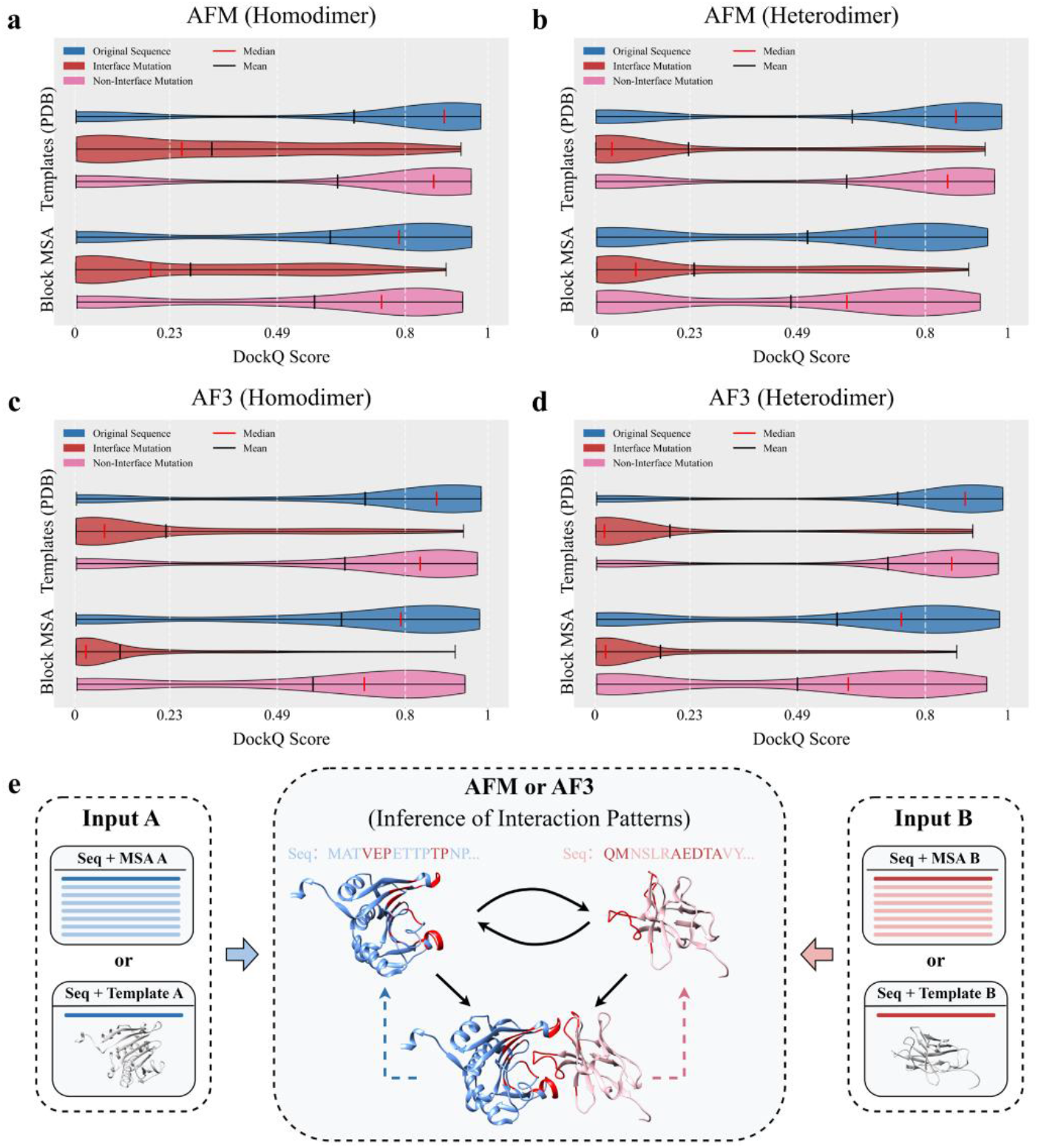
Effects of interface and non-interface mutations on complex structure prediction and the underlying mechanism. (a–d) AFM (a, b) and AF3 (c, d) performance on homodimer (a, c) and heterodimer (b, d) datasets using Block MSA or experimental determined bound monomer templates (PDB), after introducing independent random mutations at interface or non-interface residues. (e) **Modeling mechanism:** The model first establishes monomer-level geometric constraints from sequence information and MSAs or templates, and then infers compatible inter-chain interaction patterns based on the resulting monomer sequence features and structural geometry. These inter-chain constraints are subsequently iteratively refined together with intra-chain constraints, yielding a self-consistent representation for complex structure modeling.

#### 3.4 Monomer-derived features drive complex modeling

In summary, our results indicate that AFM and AF3 identify compatible interaction interfaces primarily by leveraging monomer sequences and structural geometric features (inferred from MSAs or monomer templates), rather than inter-chain coevolutionary signals (Figure 4e). This supports a mechanism in which representation-learning modules (Evoformer or Pairformer) first establish monomer-level geometric constraints, followed by the inference of inter-chain constraints. These inter- and intra-chain constraints are then iteratively refined across multiple layers and recycling rounds, enabling their joint optimization into a self-consistent complex structure.

This mechanism explains why MSA-based approaches capture interaction-induced interfacial conformational changes more effectively than template-based methods. In MSA-based inference, intra-chain constraints are progressively constructed and refined during model inference, enabling inter-chain constraints to be jointly updated in a coupled manner and thereby supporting coordinated folding and docking. In contrast, template-based inputs largely fix intra-chain geometry at an early stage by directly inheriting template-derived structures, after which inter-chain interactions are inferred primarily through geometric complementarity and sequence compatibility, resembling a more rigid docking process and limiting conformational adaptation upon binding.

It is worth noting that, in cases where strong inter-protein coevolutionary signals are present, paired MSAs may still contribute to improved prediction performance. However, our analyses demonstrate that such signals are not the dominant determinant of AlphaFold-based protein complex prediction—being virtually absent as a primary driver across almost all tested complexes. Instead, accurate complex assembly primarily emerges from the resolution of monomeric geometric constraints and their propagation toward interface formation, revealing a geometry-over-coevolution principle underlying AlphaFold-based protein complex assembly.

### 4. Validating the mechanism by tracking spatial constraints in AFM modeling

To further validate the mechanistic basis of AlphaFold-based protein complex prediction, we introduce a method termed AlphaFold-Constraint Propagation Mapping (AF-CPM) for visualizing the evolution of intra- and inter-chain geometric constraints during AFM inference. AF-CPM leverages the distogram head implemented in OpenFold^35^ to transform pair representations from successive Evoformer layers into residue–residue distance distributions. By aggregating probabilities corresponding to distances below 12 Å, we obtain residue–residue contact probability maps (Figure 5a). This framework enables direct visualization of how spatial geometric constraints are progressively inferred and propagated during complex structure prediction, providing a mechanistically interpretable view of the model’s internal inference process.

**Figure 5.**
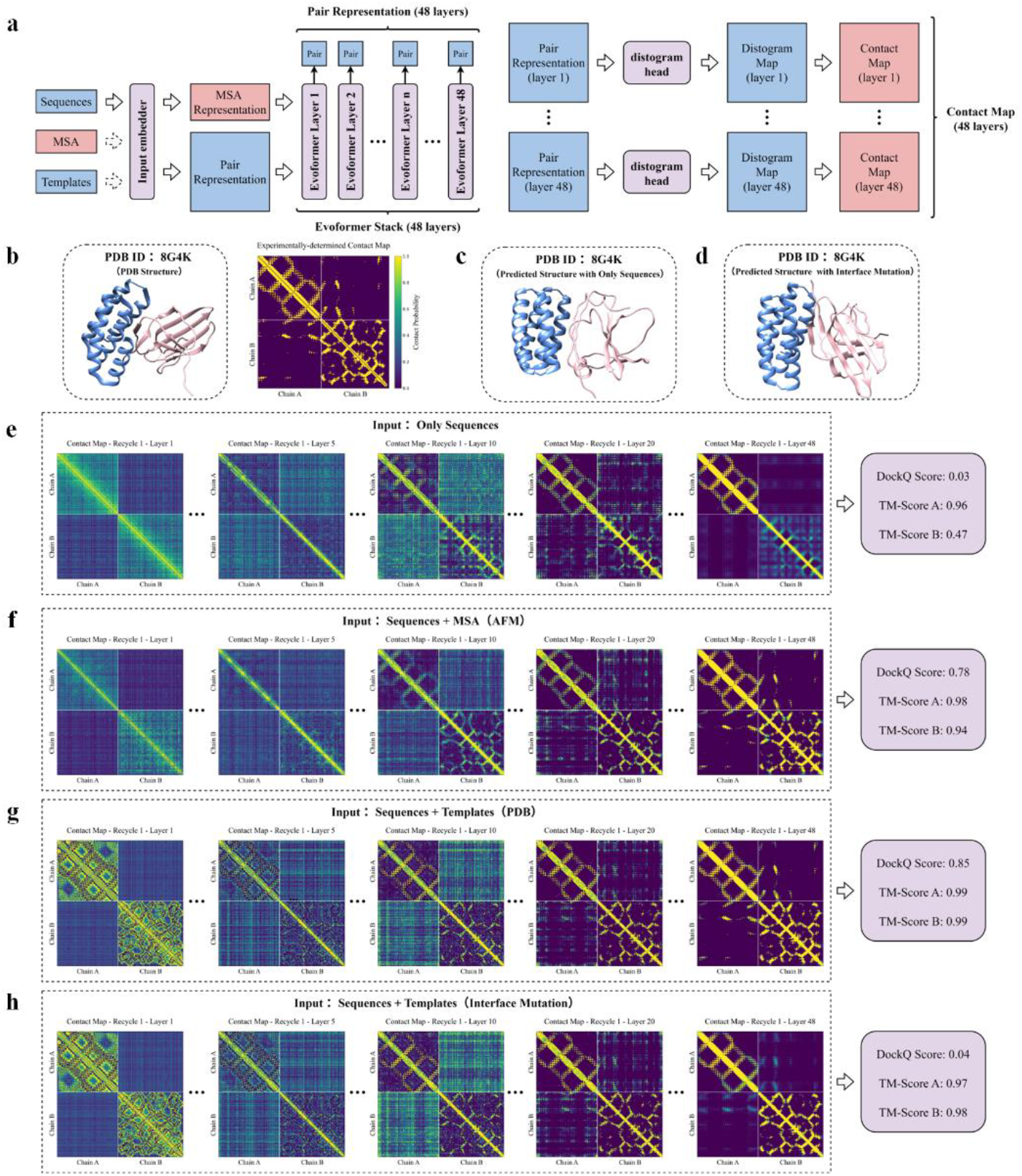
AF-CPM reveals spatial constraint propagation during AFM inference. (a) **Overview of AF-CPM:** Pair representations from each layer of the 48-layer Evoformer stack are extracted and passed to the distogram head to generate distogram maps. Contact probability maps are obtained by summing the probabilities over the first 32 distance bins of the distograms, corresponding to a distance threshold of 12 Å. These contact probability maps are computed for each layer of the Evoformer stack, yielding layer-wise representations that reflect the progressive formation of geometric constraints during complex structure prediction. (b) Experimentally determined structure of 8G4K (PDB) and its corresponding contact map derived from the experimental structure. (c) Predicted complex structure of 8G4K using only sequences. (d) Predicted complex structure of 8G4K after mutating interface residues on chain A, using mutated monomer templates. (e–h) Tracking constraint formation using AF-CPM. Evolution of spatial constraints across Evoformer layers and the corresponding prediction quality for 8G4K under different input conditions: sequence only (f), sequence + MSA (AFM) (g), sequence + templates (PDB) (h), and sequence + templates with interface mutations (i).

We applied AF-CPM to a representative heterodimeric complex (PDB ID: 8G4K) (Figure 5b). When the model was provided with protein sequences alone, without MSAs, it failed to infer intra-chain contact maps and the monomeric structure of one interacting partner (chain B), and was also unable to recover the correct inter-chain interaction interface (Figure 5c, 5e; Supplementary Figure S6). This confirms that coevolutionary information encoded in MSAs is important for accurate monomer structure inference. In contrast, when MSAs generated by AFM—including both paired and block-format alignments—were supplied, the model first established robust intra-chain contact maps and subsequently inferred inter-chain contact patterns (Figure 5f; Supplementary Figure S7). Notably, inter-chain contacts emerged only after intra-chain constraints had been established, indicating that inter-protein geometric relationships are inferred downstream of monomer-level geometry rather than being directly extracted from inter-protein coevolutionary signals.

When MSAs were omitted and experimentally determined bound-state monomer structures were provided as templates, the model accurately recovered both intra- and inter-chain contact maps and produced correct complex structures. In this template-based setting, intra-chain contact information was directly inherited from the template, bypassing the need for progressive construction through Evoformer layers, while inter-chain constraints were inferred based on the given monomeric geometry (Figure 5g; Supplementary Figure S8). This observation suggests that monomer-template–based modeling does not involve a progressive process for establishing intra-chain geometric constraints, which may explain its limited ability to capture interaction-induced conformational rearrangements at protein–protein interfaces.

To further interrogate the role of interface residues in complex assembly, we introduced random mutations at interface residues of one interaction partner (chain A) and remodeled the mutated sequence using the original monomeric structural template. Despite the preservation of strong intra-chain contact signals, inter-chain contact probabilities were almost completely abolished (Figure 5h; Supplementary Figure S9). Consequently, the model failed to recover the original binding mode, even though the monomeric structure itself remained accurately predicted (Figure 5d). These results demonstrate that accurate complex assembly cannot be achieved solely from monomer backbone geometry and highlight a critical dependence of inter-chain contact inference on interface residue identity. Together, these findings further validate the proposed protein–protein assembly mechanism in AlphaFold, whereby monomer-level geometric constraints are first established and subsequently propagated to generate cross-chain constraints through matching interface backbone geometry with residue identities, thereby enabling identification of binding modes.

### 5. Origins of limited prediction accuracy in antigen–antibody systems

Although AlphaFold models have demonstrated remarkable performance in modeling conventional protein–protein complexes, their performance on antigen–antibody systems remains substantially lower^24,25^. This limitation has often been attributed to the lack of inter-chain coevolutionary signals at the interface. However, our prior analyses of AlphaFold’s complex assembly mechanism indicate that the model primarily relies on geometric complementarity and sequence compatibility at the interface, rather than inter-chain coevolution. This raises the question of whether the absence of inter-chain coevolutionary signals is indeed the primary factor limiting performance. Motivated by this, we constructed an independent benchmark of 197 nonredundant antigen–antibody complexes to systematically reassess this assumption.

We first evaluated AFM and AF3 under multiple input conditions, including AFM MSAs, Block MSAs, and bound-state monomer templates derived from experimental structures. Across all settings, introducing inter-chain sequence pairing did not improve prediction accuracy for either antigen–antibody complexes or antibody heavy–light chain assembly. In contrast, bound-state monomer templates consistently outperformed all MSA-based approaches (Figure 6a–b). This trend is consistent with observations in conventional homo- and heterodimer systems, supporting that the reduced performance is not attributable to the absence of inter-chain coevolution, but instead reflects a geometry-driven assembly process.

**Figure 6.**
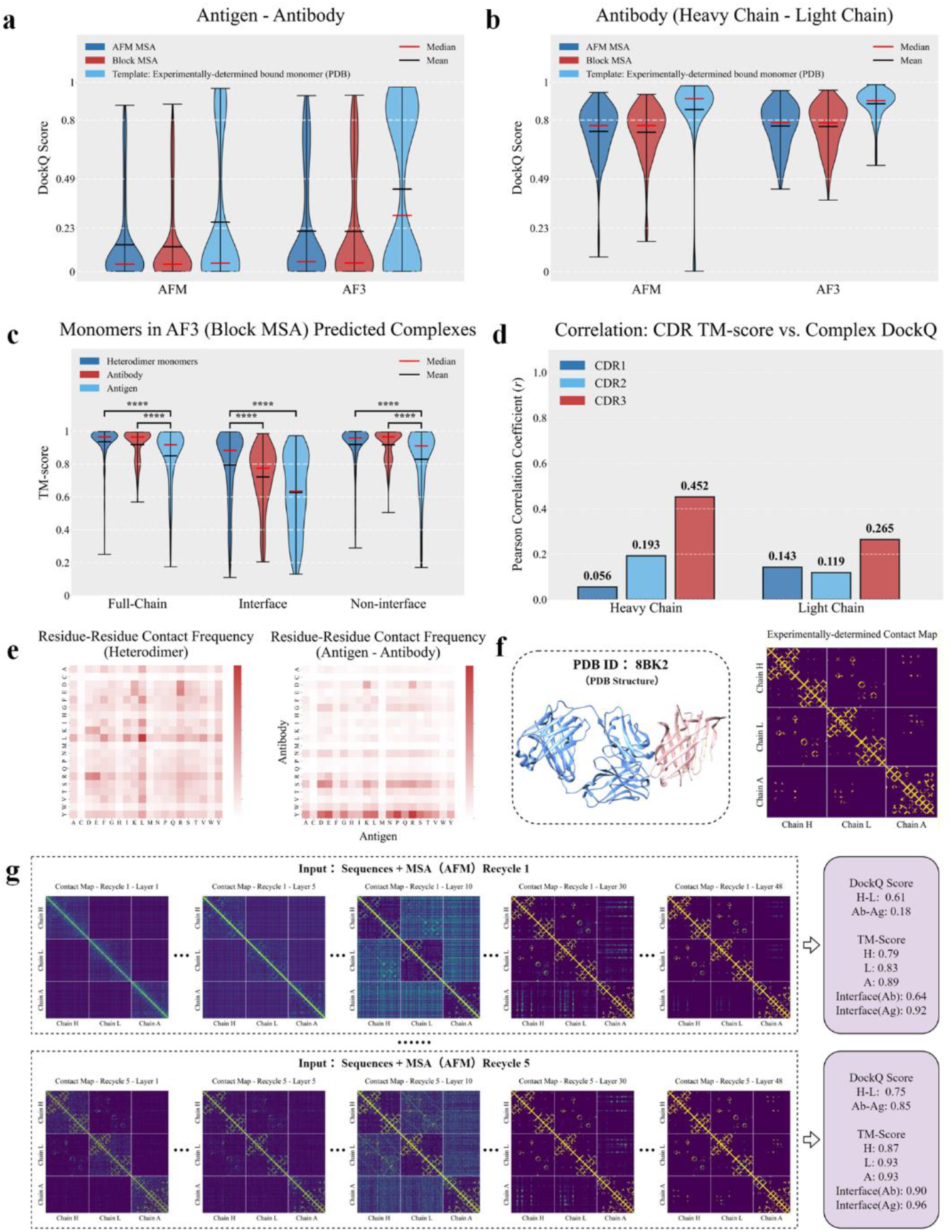
Mechanistic analysis of antigen–antibody prediction limitations. (a, b) Prediction performance of AFM and AF3 on the (a) antigen–antibody interface and (b) antibody heavy/light chain interface within the antigen–antibody dataset, evaluated under three input conditions: AFM MSA, Block MSA, and experimentally determined bound monomer templates (PDB). (c) Distribution of monomer TM-scores in AF3-predicted complex structures for heterodimers and antigen–antibody complexes at the full-chain, interface, and non-interface regions, using Block MSA as input, with interfaces defined from experimentally determined complex structures. At the full-chain and non-interface regions, antigen monomers show significantly lower accuracy than both antibody monomers and heterodimer monomers (\*\*\*\**p* < 0.0001, two-sided Mann–Whitney U test). At the interface region, both antigen and antibody monomers exhibit substantially lower accuracy than heterodimer monomers (\*\*\*\**p* < 0.0001). (d) Correlation between modeling accuracy (TM-score) of the three CDRs in antibody heavy and light chains and the prediction accuracy of the antigen–antibody interface (DockQ). (e) Statistical analysis of residue-residue contact types at heterodimer and antigen–antibody interfaces. (f) Experimental structure (left) and corresponding contact map (right) of 8BK2. (g) Evolution of constraints and predicted model accuracy for 8BK2 under AFM MSA input with different recycling rounds (1 and 5).

Nevertheless, this mechanistic insight does not fully account for the reduced prediction accuracy observed for antigen–antibody complexes. To investigate the origin of this discrepancy, we systematically compared monomer-level structural accuracy between antigen–antibody and conventional heterodimer datasets using AF3-predicted complex structures with block MSAs as input. Performance was evaluated across the full-chain, interface, and non-interface regions.

We found that antibody monomers (with the heavy and light chains merged for evaluation) show comparable full-chain and non-interface region accuracy to those of control heterodimers, whereas antigen monomer accuracy is significantly lower across full-chain, interface, and non-interface regions. More importantly, local structural accuracy at the interface—defined based on experimentally determined complex structures—on both the antigen and antibody sides is substantially lower than that of conventional heterodimers (Figure 6c). This reduced interfacial accuracy correlates with overall complex accuracy, with the most pronounced effects observed in the antibody complementarity-determining region (CDR)-H3 loop (Figure 6d; Supplementary Figure S10).

Together, these results identify a central bottleneck: the limited ability of AlphaFold to model structurally plastic interface regions. Antibody specificity is largely determined by somatically hypermutated CDRs, particularly the highly variable CDR-H3 loop, while antigen epitopes evolve under immune selection and display substantial sequence and conformational diversity. As a result, these rapidly evolving and conformationally heterogeneous interfaces constitute intrinsically challenging targets in monomeric structure prediction, thereby impairing the model’s ability to infer correct binding geometries through geometric matching.

Further statistical analysis of interfacial residue contacts revealed systematic differences between antigen–antibody complexes and conventional heterodimers. The antigen–antibody interface exhibits a distinctive residue usage pattern, with pronounced enrichment of tyrosine and serine on the antibody side—features not characteristic of typical protein–protein interfaces (Figure 6e). In addition, both heterodimer and homodimer datasets show clear differences in contact frequency distributions between successfully and unsuccessfully predicted cases (Supplementary Figure S11), highlighting a key limitation in the model’s learned interface priors and suggesting that prediction performance is governed by the alignment between interface features and learned statistical distributions. Since the model is predominantly trained on non– antigen–antibody complexes, these priors are better aligned with conventional protein– protein interaction statistics, leading to a distribution mismatch when applied to immune complexes, which may contribute as an additional source of reduced prediction performance on antigen–antibody targets.

Finally, we examined the evolution of geometric constraints in AFM predictions of antigen–antibody complexes using the AF-CPM method proposed in this study. Using PDB ID: 8BK2 as an example, AFM with its default MSA input required at least five recycling iterations to successfully assemble the complex (Figure 6f-g). In the early recycling steps, the model first establishes intra-chain constraints for each monomer, including the antibody’s heavy and light chains, as well as the antigen. It then forms inter-chain constraints between the antibody’s heavy and light chains, assembling the antibody structure correctly. However, at the end of the first recycling round, no constraints exist between the antigen and antibody. Only after five iterations of recycling do cross-chain constraints between the antibody and antigen gradually form, leading to the convergence of the complex into its correct conformation. (Figure 6g; Supplementary Figures S12–13). Notably, successful convergence coincides with improved local accuracy at the antibody interface, indicating a tight coupling between local interface refinement and global complex assembly.

Together, these findings demonstrate that the reduced accuracy of antigen–antibody prediction is not primarily attributable to the absence of inter-chain coevolution, but rather to the structural plasticity and statistically atypical nature of immune interfaces, which limit the reliable propagation of geometric constraints required for accurate interface assembly.

## Discussion

The remarkable success of AlphaFold in predicting multi-chain protein architectures has long been attributed to the direct decoding of inter-protein coevolutionary signals. In this study, we challenge the prevailing coevolution-centric view. Through systematic perturbation of model inputs and dynamic tracking of geometric constraints, we reveal a geometry-over-coevolution principle underlying AlphaFold-based protein complex assembly. We demonstrate that AlphaFold-family models do not predominantly rely on inter-protein coevolutionary signals to recognize binding partners; rather, accurate complex assembly is driven by the resolution of monomeric geometric constraints coupled with interface-specific sequence-pattern matching. This mechanistic framework elegantly reconciles the models’ high predictive performance with the widespread absence of strong inter-protein coevolutionary signals in nature.

The introduction of our AF-CPM framework provides a layer-wise view into the “black box” of AI-driven structural inference. By tracing the propagation of residue-level geometric constraints, we uncover a hierarchical assembly process in which models first establish robust intra-chain structural constraints and subsequently propagate these constraints to infer inter-chain interfaces. This layer-wise progression suggests that cross-chain organization emerges progressively through geometry refinement, rather than being primarily driven by inter-protein coevolutionary information. This interpretation is further supported by our template-based modeling results, which demonstrate that accurate monomer geometry can substantially reduce the dependence on MSA-derived evolutionary information for complex assembly. In this context, our results suggest that the principal contribution of complex MSAs may lie not in providing cross-chain evolutionary linkage per se, but in facilitating the modeling of interaction-induced conformational adaptations.

Importantly, our geometry-driven framework provides a mechanistic explanation for the persistent limitations in antigen–antibody prediction. A common explanation for this limitation has been the lack of inter-chain coevolution. However, our findings point to a different source of this difficulty: the intrinsic structural plasticity and statistical heterogeneity of immune interfaces. Unlike conventional protein–protein interactions, immune recognition surfaces exhibit pronounced local conformational variability and atypical residue compositions. These non-canonical features create a substantial distribution mismatch with the structural priors embedded in current AlphaFold models, thereby limiting the propagation of geometric constraints needed to establish accurate cross-chain interactions.

Taken together, this work provides a conceptual framework for understanding biomolecular structure prediction in the deep learning era. The geometry-over-coevolution principle shifts the focus of AI-based protein complex prediction from mining increasingly weak inter-protein coevolutionary signals in ever-larger genomic databases toward learning and propagating the geometric constraints that define molecular interfaces. This shift is particularly relevant to structurally heterogeneous, highly flexible, and non-canonical interfaces, for which accurate local geometry and sequence–geometry compatibility, rather than canonical evolutionary patterns, may ultimately determine the reliability of complex assembly.

## Materials and Methods

### 1. Dataset preparation

#### 1.1 Homodimer dataset

Crystal structures released between 1 January 2022 and 20 December 2024 were retrieved from the PDB. Structures were selected based on two criteria: resolution ≤3.0 Å and biological assemblies composed of two identical protein chains, yielding 7,620 structures. To remove sequence redundancy, the remaining structures were clustered using MMseqs2^36^ at a 40% sequence identity threshold, resulting in 2,509 clusters. From these, 200 homodimer complexes were randomly selected to construct a manageable benchmark dataset, each representing a distinct cluster and restricted to entries with a unique biological assembly.

#### 1.2 Heterodimer dataset

Using the same temporal and resolution criteria as for the homodimer dataset, 2,310 PDB entries containing heterodimeric structures were retrieved from the PDB. To remove sequence redundancy, heterodimeric chains were clustered using MMseqs2 at a 40% sequence identity threshold, resulting in 691 clusters. Each heterodimeric chain pair was then assigned a cluster pair identifier based on the cluster membership of its constituent chains, treating the pair as unordered (i.e., chain cluster A–chain cluster B and chain cluster B–chain cluster A were considered equivalent), and pairs in which both chains belonged to the same cluster were excluded to avoid sequence homology between interacting partners. Heterodimer complexes sharing the same cluster pair identifier were grouped, and one representative complex was randomly selected from each group, restricted to entries with a unique biological assembly, yielding 317 non-redundant heterodimer complexes. One covalently linked dimer (PDB ID: 8DLE) was subsequently excluded, resulting in a final benchmark set of 316 heterodimer complexes.

#### 1.3 Antigen–antibody dataset

Antigen–antibody complex structures were retrieved from the Structural Antibody Database^37^ (SAbDab) using the “search structures by attribute” function. Structures were filtered with a resolution cutoff of ≤4.0 Å and antigen type restricted to proteins. To avoid overlap with the training data of AFM/AF3, only structures released between 1 January 2022 and 18 March 2024 were included, yielding 2083 antigen–antibody complexes. To remove redundancy, antigen sequences were clustered using MMseqs2 at a 40% sequence identity threshold. Antigen–antibody complexes sharing the same antigen identifier were grouped, and one representative complex was randomly selected from each group, resulting in a final test set of 197 antigen–antibody complexes.

### 2. Structure prediction implementation

#### 2.1 AFM

Protein complex structure prediction was performed using AlphaFold2_multimer_v3 as implemented in LocalColabFold (https://github.com/YoshitakaMo/localcolabfold). For each target, five models were generated using the five sets of model parameters with a fixed random seed (0). The model with the highest ranking score was selected as the final prediction.

#### 2.2 AF3

Protein complex structure prediction was performed using the official AlphaFold3 scripts (https://github.com/google-deepmind/alphafold3). For each target, five models were generated with a fixed random seed (1). The model with the highest ranking score was selected as the final prediction.

### 3. Design and implementation of interface and non-interface mutations

#### 3.1 Selection of mutation sites

Protein–protein interaction interfaces were defined from dimeric complex structures retrieved from the PDB. A residue was classified as an interface residue if any of its heavy atoms lay within 5 Å of any heavy atom from the partner chain, whereas residues with a minimum heavy-atom distance greater than 8 Å from all partner-chain atoms were defined as non-interface residues.

For heterodimers, mutations were introduced randomly in a single chain. For homodimers, to preserve structural symmetry, only residues with consistent interface or non-interface classification at equivalent positions in both chains were considered, and mutations were introduced at these positions in both chains simultaneously. Five homodimers lacking residues that satisfied these criteria were excluded from further analysis.

Interface and non-interface perturbations were performed in separate experiments. For each category, the number of mutated residues was limited to at most 10% of the chain length; when the number of eligible residues exceeded this threshold, a random subset of residues corresponding to 10% of the chain length was selected. All selected residues were mutated to glycine to standardize the perturbation.

#### 3.2 Generation of MSAs for mutated sequences

The mutated sequences were submitted to LocalColabFold to generate MSAs using the AlphaFold2_multimer_v3 model. MSAs were preprocessed to remove lowercase insertion characters, and the paired segment was extracted and converted into a block MSA format for subsequent complex structure prediction.

#### 3.3 Generation of monomer templates for mutated sequences

Single-chain templates were obtained from experimentally determined bound-state structures in the PDB. Mutated sequences were aligned to these templates, and homology modeling was performed using MODELLER^38^ to incorporate mutations while preserving the backbone conformation. The resulting monomer templates were used for complex structure prediction with the mutated sequences.

### 4. Quantitative assessment of predictions

#### 4.1 Protein–protein complex structure assessment

The quality of protein–protein complex structure predictions was evaluated using DockQ (https://github.com/wallnerlab/DockQ).The prediction quality is categorized based on the DockQ score as follows: 0 ∼ 0.23 as incorrect, 0.23 ∼ 0.49 as acceptable quality, 0.49 ∼ 0.8 as medium quality, and 0.8 ∼ 1 as high quality. For mutation experiments, to address sequence–structure mismatches introduced by mutations, predicted structures were remodeled onto the corresponding wild-type sequences using MODELLER prior to DockQ calculation. For antigen–antibody complexes, which involve three chains (antibody heavy and light chains and one antigen chain), DockQ was computed under two evaluation settings. For antigen–antibody interface evaluation, the antibody heavy and light chains were merged into a single chain, whereas for evaluation of the antibody heavy–light chain interface, the two chains were treated as separate entities.

#### 4.2 Monomer structure assessment

The accuracy of monomer structures was assessed using TM-score, computed with USalign^39^. The multimeric alignment option (-mm) was set to 0 to perform pairwise monomer structure alignment. To compare interface and non-interface regions, the interface region was defined as residues within 5 Å of any heavy atom on the partner chain, whereas the non-interface region comprised residues with a minimum distance greater than 8 Å to any heavy atom on the partner chain. For antibody structures, the heavy and light chains were treated as a single entity and evaluated jointly, except when analyzing CDR loops (Figure 6d). Accordingly, antibody monomer accuracy in Figure 6c and Supplementary Figure S10a was computed based on the combined heavy–light chain structure.

#### 4.3 Statistical analysis

Because DockQ and TM-score distributions deviated from normality, statistical comparisons between groups were performed using the two-sided Mann–Whitney U test^40^. Differences with *p*< 0.05 were considered statistically significant.

### 5. Implementation of AF-CPM for tracking geometric constraints

We employed OpenFold (v2.2.0) with pretrained AFM parameters (params_model_3_multimer_v3) to track the propagation of residue-level geometric constraints during protein complex prediction. The OpenFold pipeline was modified to support customized template and MSA inputs.

To extract intermediate structural representations, we further modified OpenFold to retrieve pair representations from each of the 48 layers of the Evoformer stack, which were subsequently converted into residue–residue distance distributions using the Distogram head. The resulting distance distributions were normalized using a softmax function, such that each bin represents the predicted probability that the Cβ atoms of a residue pair fall within the corresponding distance interval. Contact probability maps were then derived by summing the probabilities of the first 32 distance bins, corresponding to a distance threshold of 12 Å. The resulting layer-wise contact probability maps capture the progressive establishment of geometric constraints during complex structure prediction.

## Author Contributions

C.Y. conceived the project, designed the experiments, and supervised the study. S.L. performed the experiments. S.L., Z.M., and C.Y. analyzed the data. S.L. and C.Y. drafted the manuscript. All authors discussed the results and revised the manuscript.

## Data, Code, and Materials Availability

The data supporting the findings of this study, including the PDB IDs of all datasets used and the raw data underlying the main results, are available at https://github.com/ChengfeiYan/AF-CPM/tree/main/data. The PDB structures explicitly referenced in the manuscript are PDB ID 8BK2 and PDB ID 8G4K; their corresponding PDB validation reports are provided as supplementary materials. The code implementing AF-CPM is available in the same repository. Additional scripts for data processing and analysis are available from the corresponding author upon reasonable request.

## Acknowledgements

The work was supported by the National Natural Science Foundation of China (grant number: 32571438 and 32101001). Computational resources were provided by the HPC platform of Huazhong University of Science and Technology.

## Supplementary Materials

### Supplementary Text S1

#### Structural similarity filtering using Foldseek-Multimer

To control for potential confounding effects arising from structural similarity between the time-partitioned benchmark datasets and the AlphaFold training set, we used Foldseek-Multimer^41^ to identify structurally similar complexes. Each query complex from the homodimer and heterodimer datasets was searched against the PDB database, restricted to entries deposited before 2021-09-30.

Only hits in which both chains of the query dimer aligned to the same PDB structure were considered valid. For each hit, the complexqtmscore provided by Foldseek-Multimer was recorded, and the maximum value across all hits was taken as the final similarity score for each query complex. For query complexes with no valid hits, the similarity score was set to 0.

Based on this criterion, we defined low structural similarity subsets as follows: 43 homodimer complexes with a similarity score < 0.6 and 66 heterodimer complexes with a similarity score < 0.4.

### Supplementary Figures

**Figure S1.**
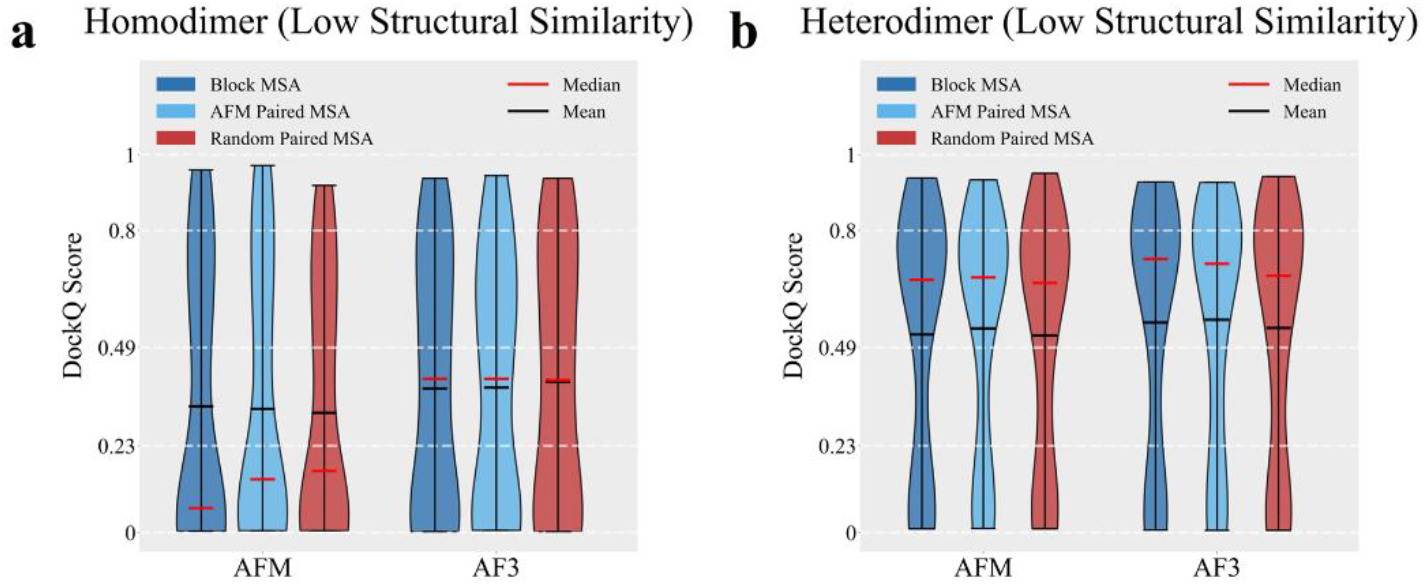
Comparison of AFM and AF3 complex prediction performance across different MSA input types on low-similarity subsets. Control for structural similarity to the AlphaFold training set. Complexes with high structural similarity to pre-2021 PDB entries were identified using Foldseek-Multimer and removed to construct low-similarity subsets for homodimer (a) and heterodimer benchmarks (b). Analyses on these filtered subsets yielded results consistent with those from the full datasets, indicating no significant differences (*p*>0.05, two-sided Mann–Whitney U test) among the three MSA input types. See Supplementary Text S1 for details.

**Figure S2.**
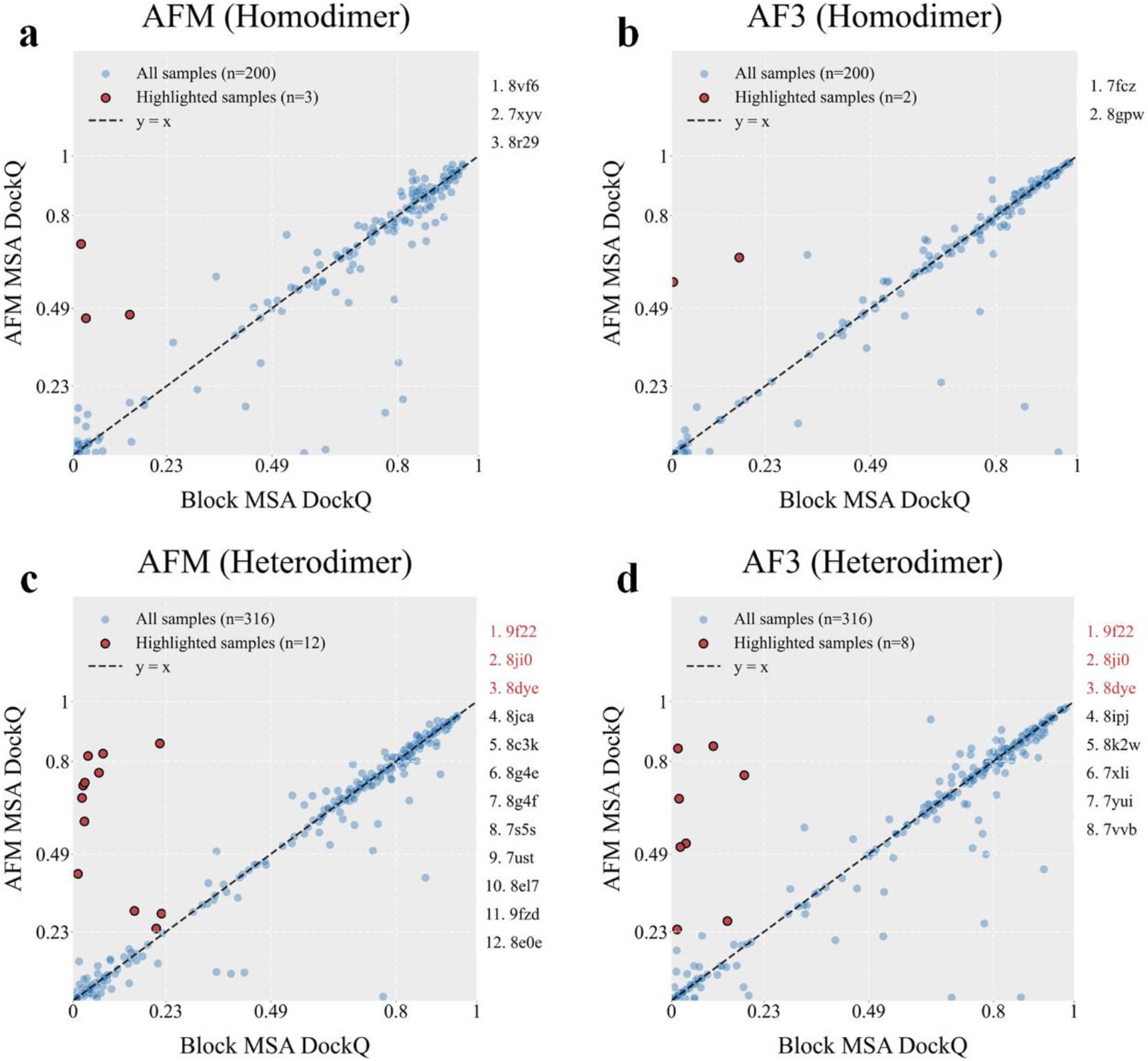
DockQ scores for complexes predicted by AFM (a, c) and AF3 (b, d) using Block MSA versus AFM MSA, on homodimer (a, b) and heterodimer (c, d) datasets. Red-highlighted points represent samples with successful predictions (DockQ ≥ 0.23) by AFM MSA but unsuccessful predictions (DockQ < 0.23) by Block MSA. Corresponding PDB IDs are listed on the right. In the homodimer dataset, no overlap is observed between AFM and AF3 predictions. In the heterodimer dataset, only three samples overlap between AFM and AF3 predictions, with their PDB IDs also highlighted in red.

**Figure S3.**
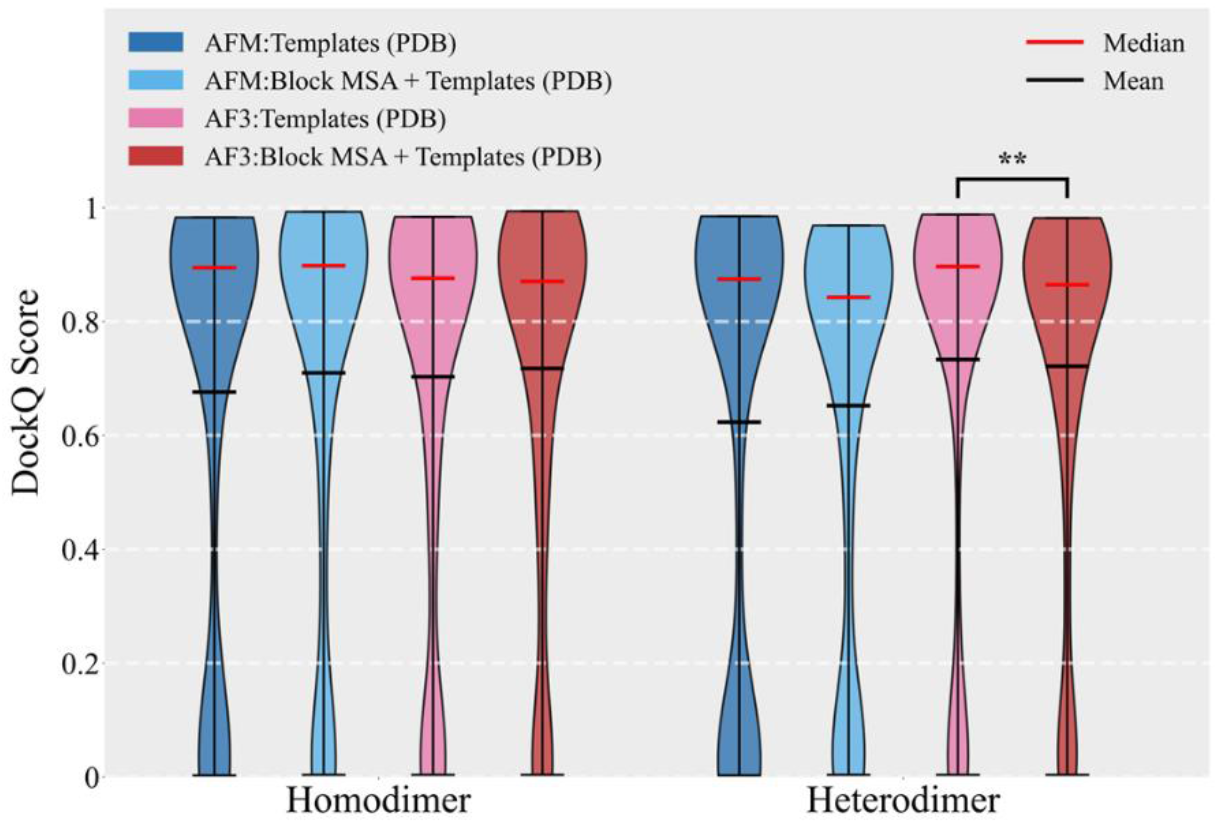
Comparison of predictions using experimental bound templates with or without Block MSA. On the homodimer dataset, no significant difference in performance is observed between the two conditions (*p* > 0.05, two-sided Mann–Whitney U test). On the heterodimer dataset, the effect of including Block MSA differs between models: for AF3, it results in a slight but significant decrease in performance (\*\**p* < 0.01), whereas for AFM, no significant difference is observed (*p* > 0.05).

**Figure S4.**
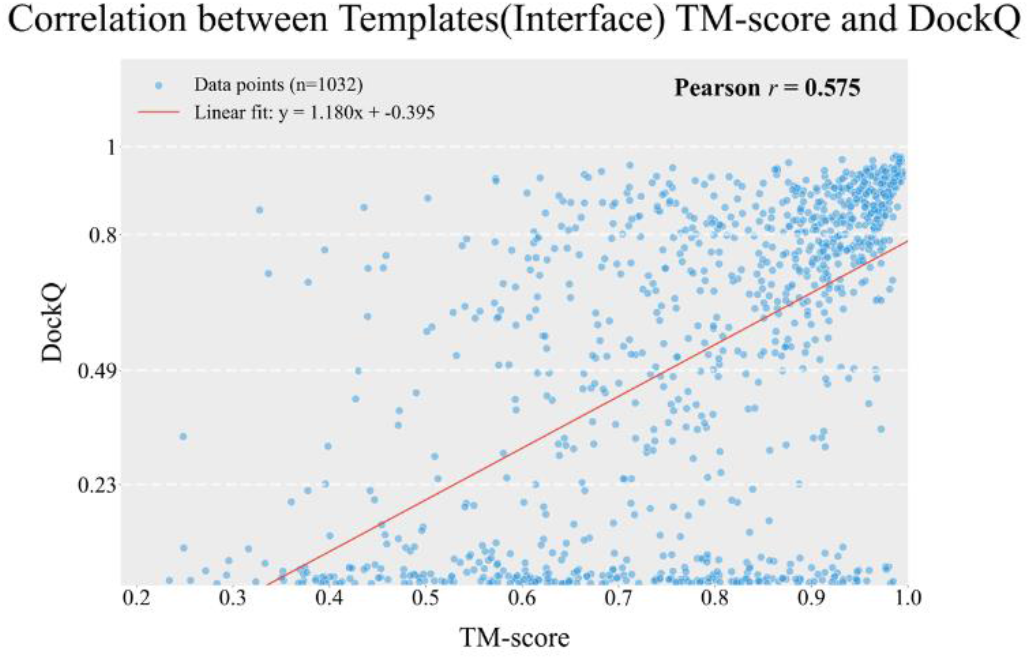
Relationship between monomer template interface accuracy and complex prediction accuracy. Monomer template interface accuracy, measured as TM-score relative to experimentally determined bound-state monomers, is correlated with DockQ across homomeric and heteromeric datasets (Pearson *r*=0.575).

**Figure S5.**
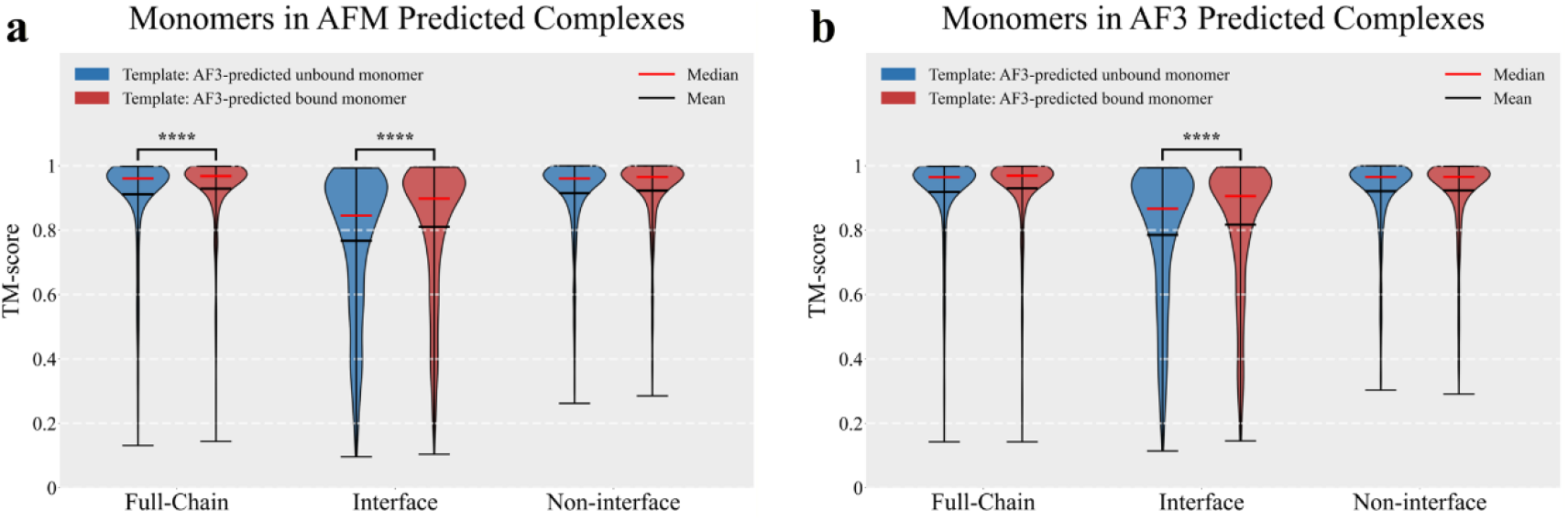
Effect of AF3-predicted bound and unbound monomer templates on monomer accuracy in complex predictions. (a) TM-score distributions of monomers in AFM-predicted complexes, comparing structures derived from AF3-predicted bound versus unbound monomer templates against experimentally determined bound monomers (PDB), evaluated at the full-chain, interface, and non-interface regions. Significant differences were observed at the full-chain and interface regions (\*\*\*\**p* < 0.0001, two-sided Mann–Whitney U test). (b) Same as in (a), but for AF3-predicted complexes. Significant differences were only observed at the interface regions (\*\*\*\**p* < 0.0001).

**Figure S6.**
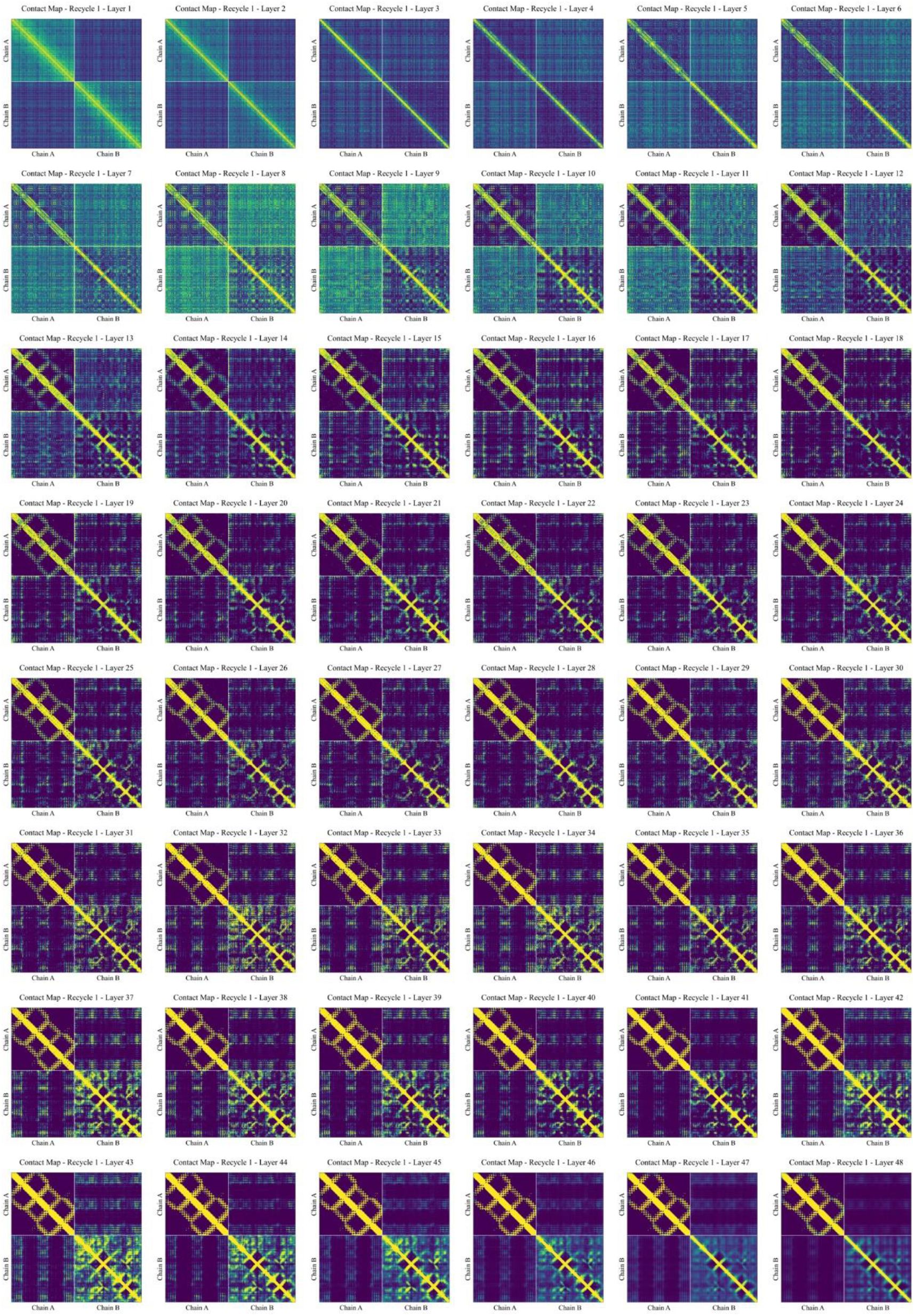
Spatial constraint propagation tracked by AF-CPM across the 48 Evoformer layers during the first recycling round for sample 8G4K with sequence-only input.

**Figure S7.**
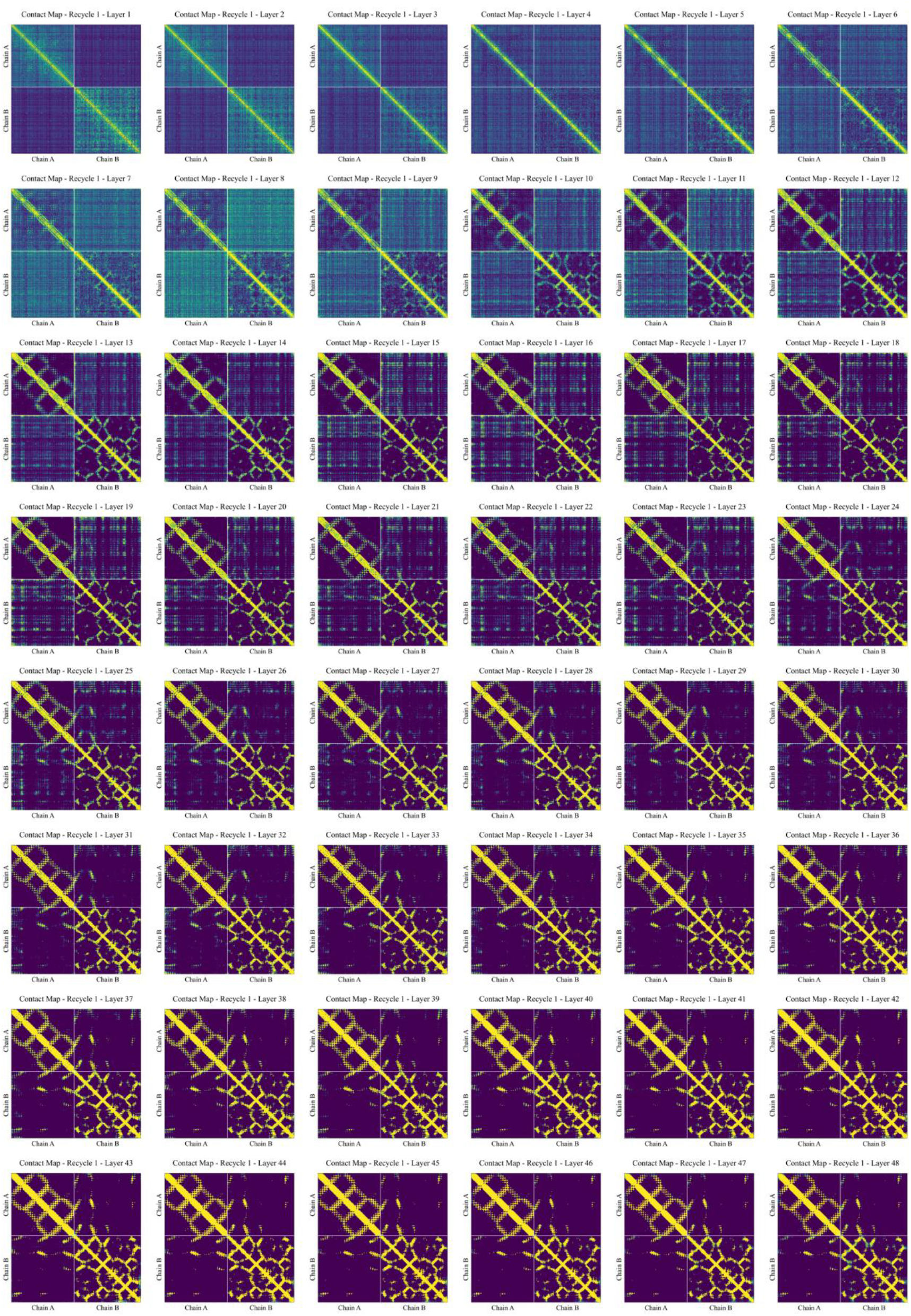
Spatial constraint propagation tracked by AF-CPM across the 48 Evoformer layers during the first recycling round for sample 8G4K with sequence + MSA input.

**Figure S8.**
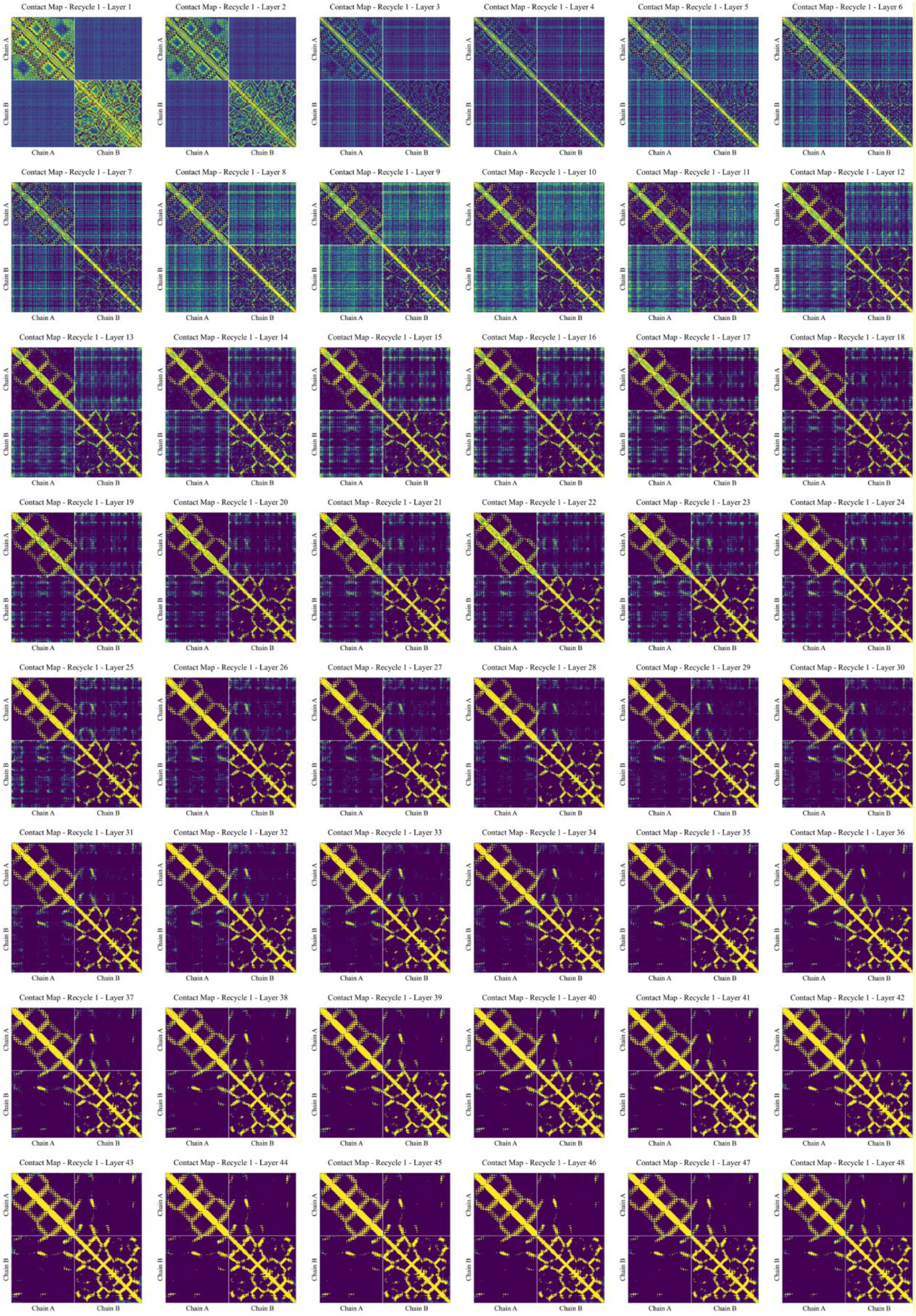
Spatial constraint propagation tracked by AF-CPM across the 48 Evoformer layers during the first recycling round for sample 8G4K with sequence + experimental bound templates (PDB) input.

**Figure S9.**
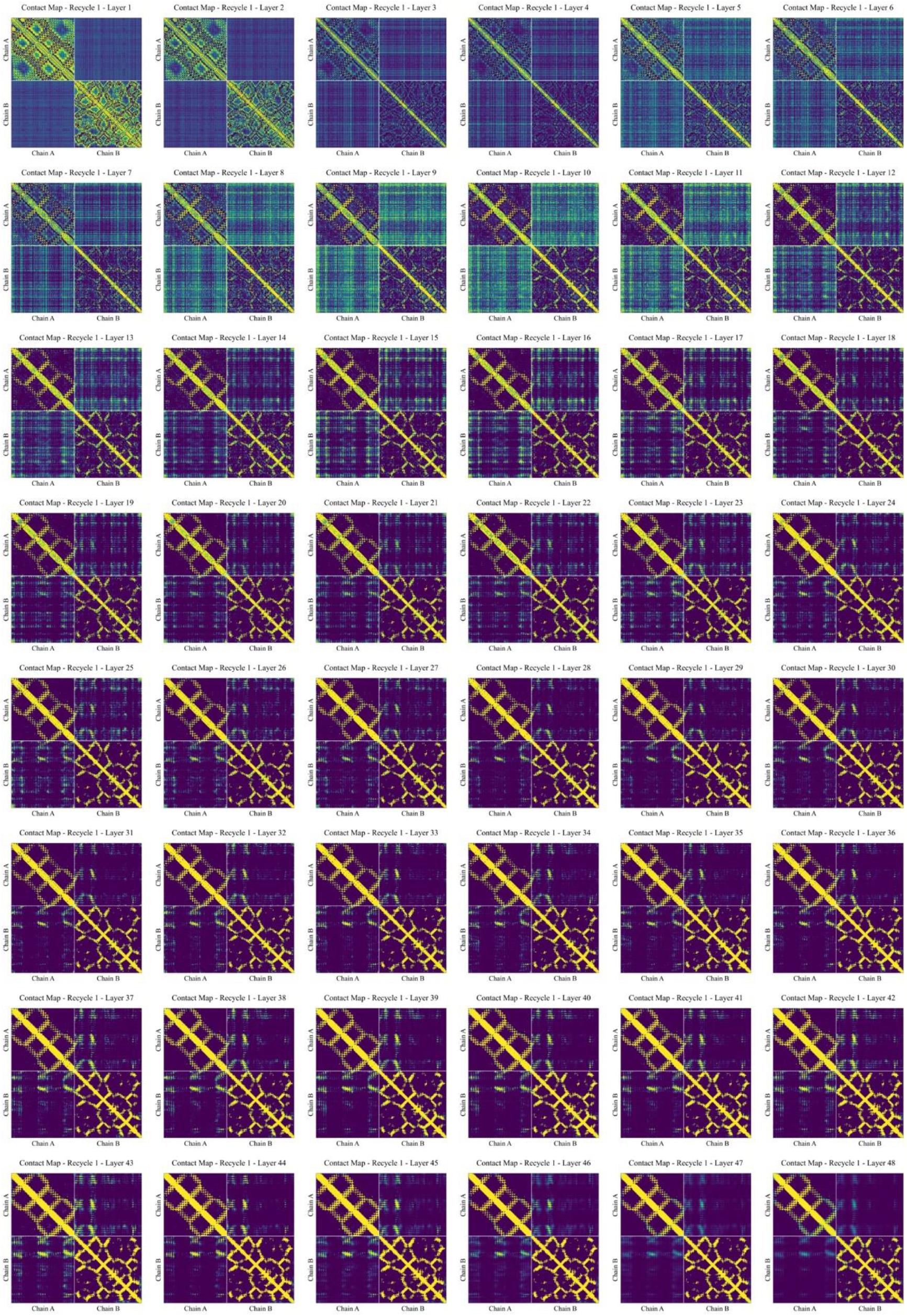
Spatial constraint propagation tracked by AF-CPM across the 48 Evoformer layers during the first recycling round for sample 8G4K with sequence + templates (interface mutation) input.

**Figure S10.**
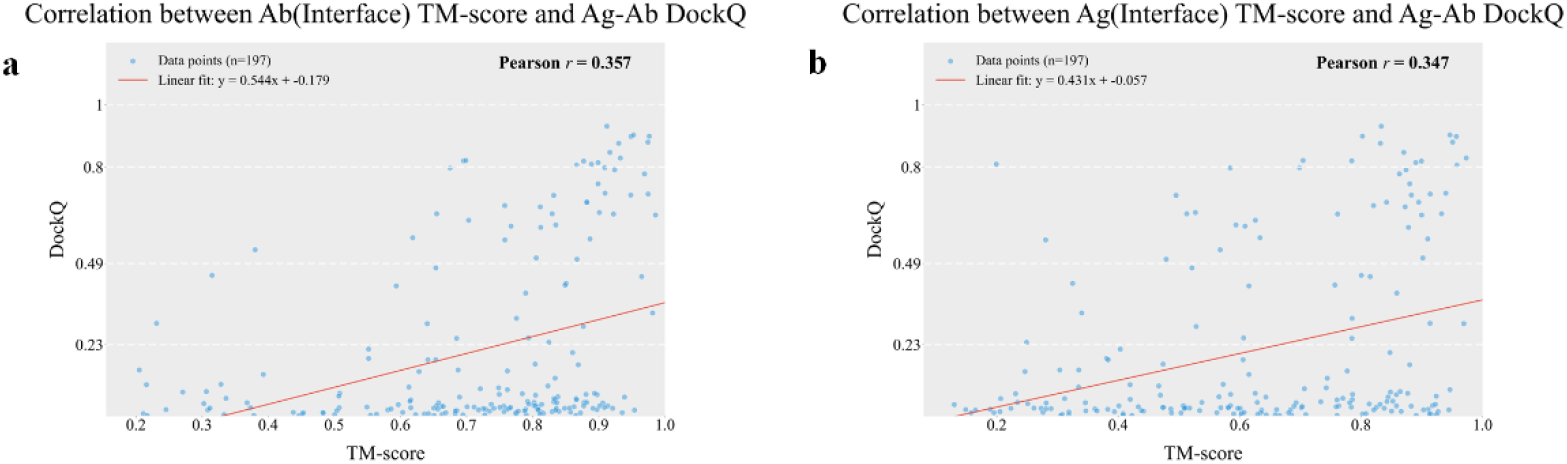
Correlation between local interface structural accuracy and complex structure prediction accuracy in antigen–antibody complexes. (a) The correlation between the TM-score of the antibody interface region and the DockQ score of the antigen–antibody complex structure. (b) The correlation between the TM-score of the antigen interface region and the DockQ score of the antigen–antibody complex structure.

**Figure S11.**
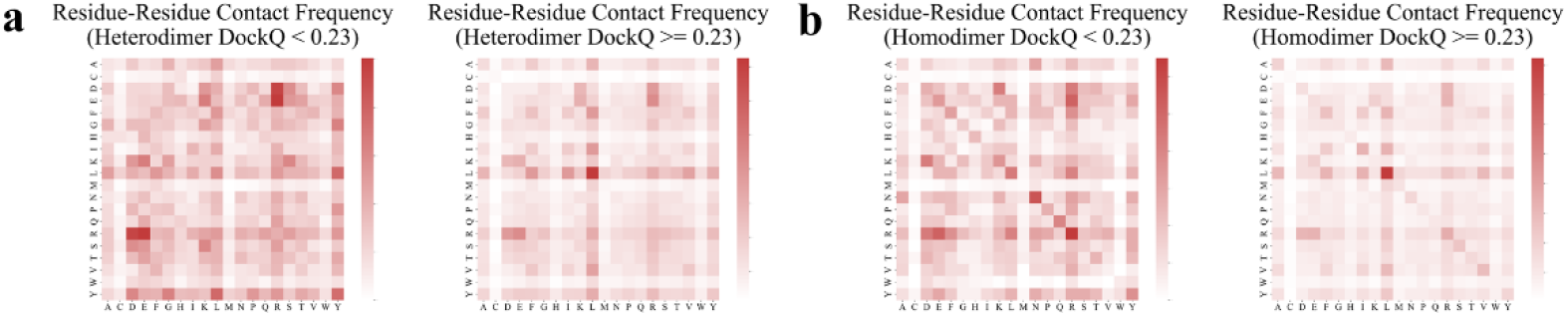
Statistics of residue-residue contact types in heterodimer and homodimer datasets. Residue-residue contact types were analyzed for successful and failed predictions in the heterodimer (a) and homodimer (b) datasets using AF3 with sequence + templates (PDB) input.

**Figure S12.**
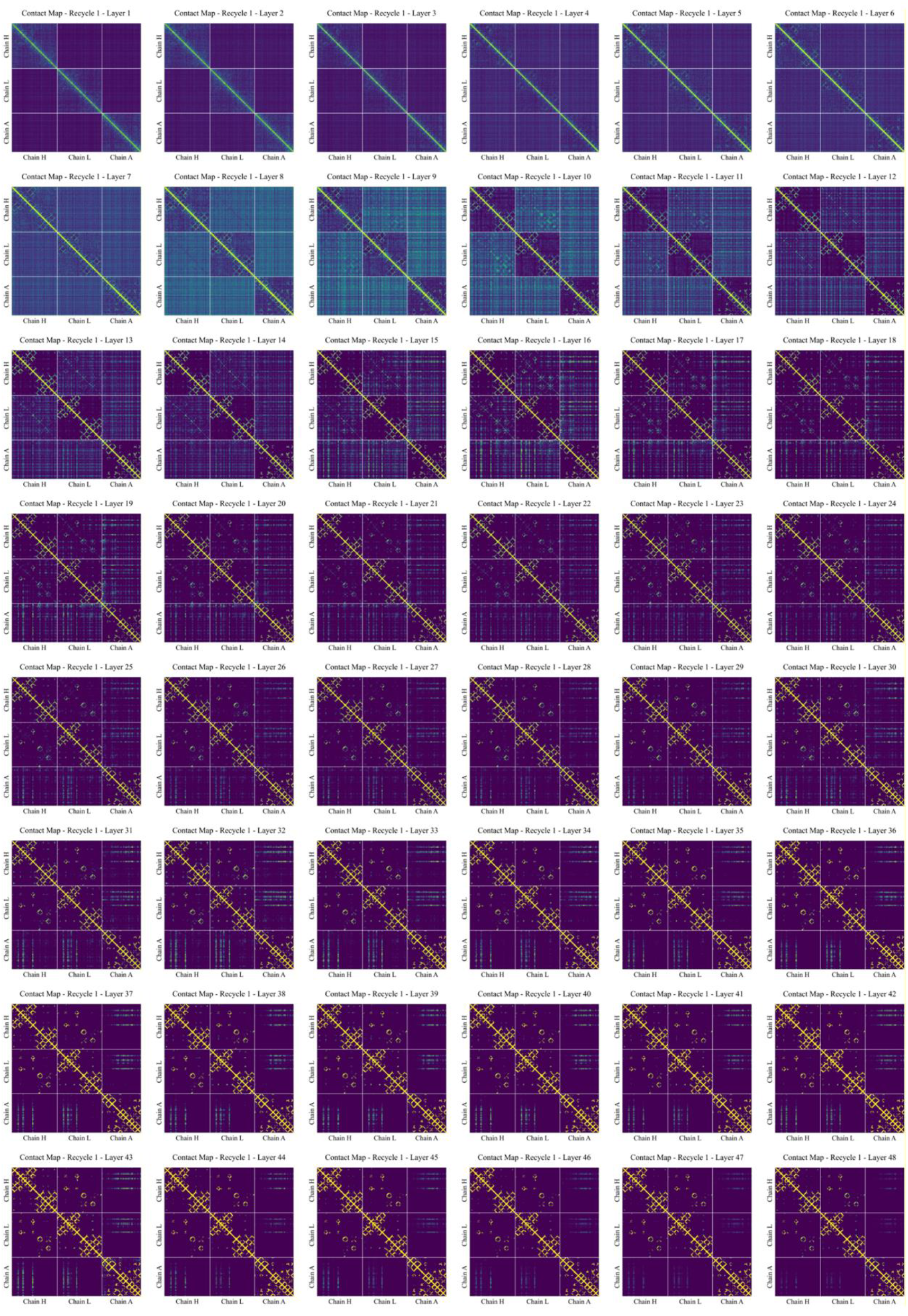
Spatial constraint propagation tracked by AF-CPM across the 48 Evoformer layers during the first recycling round for sample 8BK2 with sequence + MSA (AFM) input.

**Figure S13.**
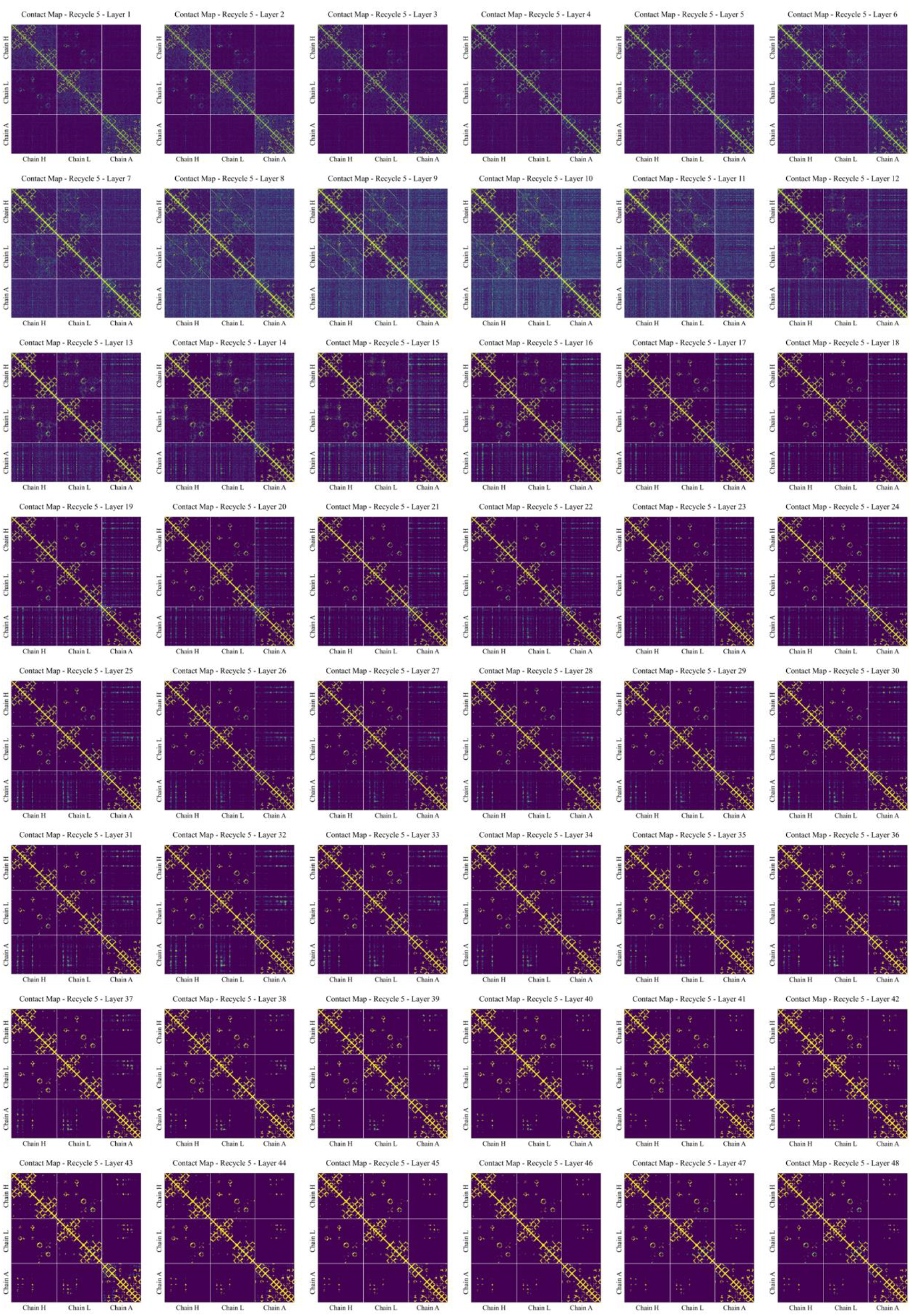
Spatial constraint propagation tracked by AF-CPM across the 48 Evoformer layers during the fifth recycling round for sample 8BK2 with sequence + MSA (AFM) input.

